# StemBond hydrogels optimise the mechanical microenvironment for embryonic stem cells

**DOI:** 10.1101/768762

**Authors:** Céline Labouesse, Chibeza C. Agley, Bao Xiu Tan, Moritz Hofer, Alex Winkel, Giuliano G. Stirparo, Hannah T. Stuart, Christophe M. Verstreken, William Mansfield, Paul Bertone, Kristian Franze, Jose C. R. Silva, Kevin J. Chalut

## Abstract

Studies of mechanical signalling are typically performed by comparing cells cultured on soft and stiff hydrogel-based substrates. However, it is challenging to independently and robustly control both substrate stiffness and tethering of extracellular matrix (ECM) to substrates, making ECM tethering a potentially confounding variable in mechanical signalling investigations. Moreover, poor ECM tethering can lead to weak cell attachment. To address this, we developed StemBond hydrogels, a hydrogel formulation in which ECM tethering is stable and can be varied independently of stiffness. We show that soft StemBond hydrogels provide an optimal format for culturing embryonic stem (ES) cells. We find that soft StemBond substrates improve the homogeneity of ES cell populations, boost their self-renewal, and increase the efficiency of cellular reprogramming. Our findings underline how soft microenvironments impact mechanosensitive signalling pathways regulating self-renewal and differentiation, indicating that optimising the complete mechanical microenvironment will offer greater control over stem cell fate specification.

## INTRODUCTION

The physical properties of the cellular microenvironment, such as substrate stiffness and adhesiveness, have a strong influence on stem cell function, including maintenance and differentiation (Crowder, Leonardo, Whittaker, Papathanasiou, & Stevens, 2016; Lutolf, Gilbert, & Blau, 2009; Sun, Chen, & Fu, 2012). To control substrate stiffness *in vitro*, polyacrylamide (PAAm) hydrogels have been used extensively because their stiffness can be varied over several orders of magnitude within the physiological range, and their surface can be functionalised by the tethering of extra-cellular matrix (ECM) proteins. In one of the most widely used approaches in the field, polymerised PAAm hydrogels are treated with a hetero-bifunctional crosslinker, the most commonly employed one being sulfo-SANPAH, though alternative methods also exist (Damljanović, Lagerholm, & Jacobson, 2005; Grevesse, Versaevel, Circelli, Desprez, & Gabriele, 2013; Kandow, Georges, Janmey, & Beningo, 2007). Sulfo-SANPAH-treated PAAm hydrogels have been successfully used, for example, to study the impact of substrate stiffness on regulating fate choices of mesenchymal stem cells (Engler, Sen, Sweeney, & Discher, 2006; Park et al., 2011). There have also been a handful of studies using these substrates to culture cells from soft tissue, such as embryonic stem (ES) cells (Chowdhury, Li, et al., 2010). However, as we show here, cells – such as ES cells – that do not develop strong focal adhesions (Xia, Yim, & Kanchanawong, 2019) detach from the substrates after a short time in culture. This limitation of commonly used PAAm substrates represents a significant bottleneck for studies in stem cell mechanobiology.

The detachment of cells from PAAm substrates could be due to a loss of ECM-substrate coupling, which is mediated by the reaction between the nitrophenyl azide group of Sulfo-SANPAH and polyacrylamide upon photo-activation. The non-specific nature of this reaction and the highly reactive nature of the photoactivated intermediate species render the binding to the substrate highly variable and difficult to control. Furthermore, it has been suggested that the effective crosslinker density depends not only on sulfo-SANPAH concentration, but also on hydrogel pore size (Trappmann et al., 2012). The importance of pore size has been debated given that adult stem cells grown on substrates above 4 kPa are functionally insensitive to changes in ECM tethering density (Wen et al., 2014). However, pore size could become relevant when culturing stem cells on much softer substrates below 1kPa, which possess larger pores, as is typically done with cells from soft tissue such as embryonic or neural tissue (Barriga, Franze, Charras, & Mayor, 2018; Segel et al., 2019).

To facilitate long-time culture of ES cells on PAAm substrates of varying stiffness, we here sought an alternative approach to PAAm functionalisation that would allow us to better control ECM tethering in a physiological range of hydrogel stiffness. We chose a method that would allow us to easily regulate ECM tethering by incorporating a co-polymer in the PAAm that can be subsequently covalently bound to ECM proteins (Pless, Lee, Roseman, & Schnaar, 1983). The ECM tethering density is here controlled by the concentration of co-polymer, and the stability of the functionalisation is ensured by the specificity of the polymerisation reaction. We have thus developed a range of substrates, which we call ‘StemBond’ hydrogels, with control of both ECM tethering density and stiffness for stem cell culture and investigations of mechanical signalling.

We compare StemBond hydrogels to sulfo-SANPAH treated PAAm to assess their effectiveness in the maintenance of ES cell self-renewal in multiple culture conditions. ES cells are uniquely characterised by their pluripotency, and their ability to self-renew in culture when in the presence of the appropriate soluble signals regulating key pluripotency pathways. They can be propagated as a homogeneous population in a naïve, *i.e*. fully uncommitted, state by the dual inhibition (“2i”) of the GSK3β and the MEK/ERK pathways (Wray, Kalkan, & Smith, 2010; Ying et al., 2008). Alternatively, a heterogeneous population with a mix of naïve and more differentiated cells can be maintained in serum-containing medium supplemented with the cytokine Leukemia Inhibitory Factor (LIF), an activator of JAK/STAT pathway (Serum+LIF medium or “S+L”) (Matsuda et al., 1999; Niwa, Burdon, Chambers, & Smith, 1998; Toyooka, Shimosato, Murakami, Takahashi, & Niwa, 2008). Interestingly, all of the three aforementioned pathways – GSK3β, MEK/ERK and JAK/STAT – have been shown to be mechanosensitive in other cell systems (Fernández-Sánchez et al., 2015; Lammerding et al., 2004; Li et al., 2010; Paszek et al., 2005). Moreover, a few studies have suggested that ES cells could sense and respond to substrate adhesiveness (Murray et al., 2013) and stiffness (Chowdhury, Li, et al., 2010; Chowdhury, Na, et al., 2010; Evans et al., 2009). However, these studies were performed either on plastic substrates or on sulfo-SANPAH functionalised soft substrates, neither of which offered good control of both stiffness and ECM tethering. As a result, exactly how the mechanical microenvironment impacts the pathways regulating ES cell self-renewal or loss of pluripotency still remains to be elucidated.

Using our StemBond hydrogels, we found that substrate stiffness highly impacts the maintenance of naïve pluripotency in ES cells, both transcriptionally and functionally. We also found a modest impact of ECM tethering on ES cell function. Even in minimal medium conditions (*i.e.* a single chemical inhibitor), in which ES cells normally differentiate, self-renewal was supported by soft substrates. We additionally found that the efficiency of reprogramming into naïve pluripotency was enhanced on soft substrates. Finally, we showed that the functional boost on soft substrates is mediated by stiffness-dependent JAK/STAT3 and MEK/ERK signalling.

## RESULTS

### StemBond hydrogels have controlled stiffness and ECM tethering density

In order to establish an optimal mechanical microenvironment for ES cells, and to enable their stable substrate attachment, we developed a new protocol for the fabrication of PAAm hydrogels. We started from a precursor solution for a standard PAAm hydrogel below 1kPa, approaching the low stiffness of the pre-implantation embryo from which ES cells are sourced (varying between 300Pa to 1000Pa according to unpublished AFM data from our group). We added to the precursor solution the co-factor 6-Acrylamidohexanoic acid (AHA), which can bind to acrylamide chains without crosslinking them (Yip et al., 2013). The terminal carboxyl groups of the AHA serve as anchorage points for covalent ECM protein binding by first forming an amine-reactive ester through a carbodiimide reaction. As with the widely used sulfo-SANPAH, this ester can react with any primary amine of, for example, lysine chains, to tether ECM proteins to the surface (**Figure 1A**). However, compared to the sulfo-SANPAH method, this method allows better control over the binding and surface density of the carboxyl groups because the AHA chains co-polymerise with acrylamide. We thus varied AHA concentration to have various levels of ECM tethering density (**Figure 1B**). We adapted the ratios of acrylamide to bis-acrylamide to ensure a constant stiffness over the range of AHA concentrations (**Figure 1C**).

**Figure 1:**
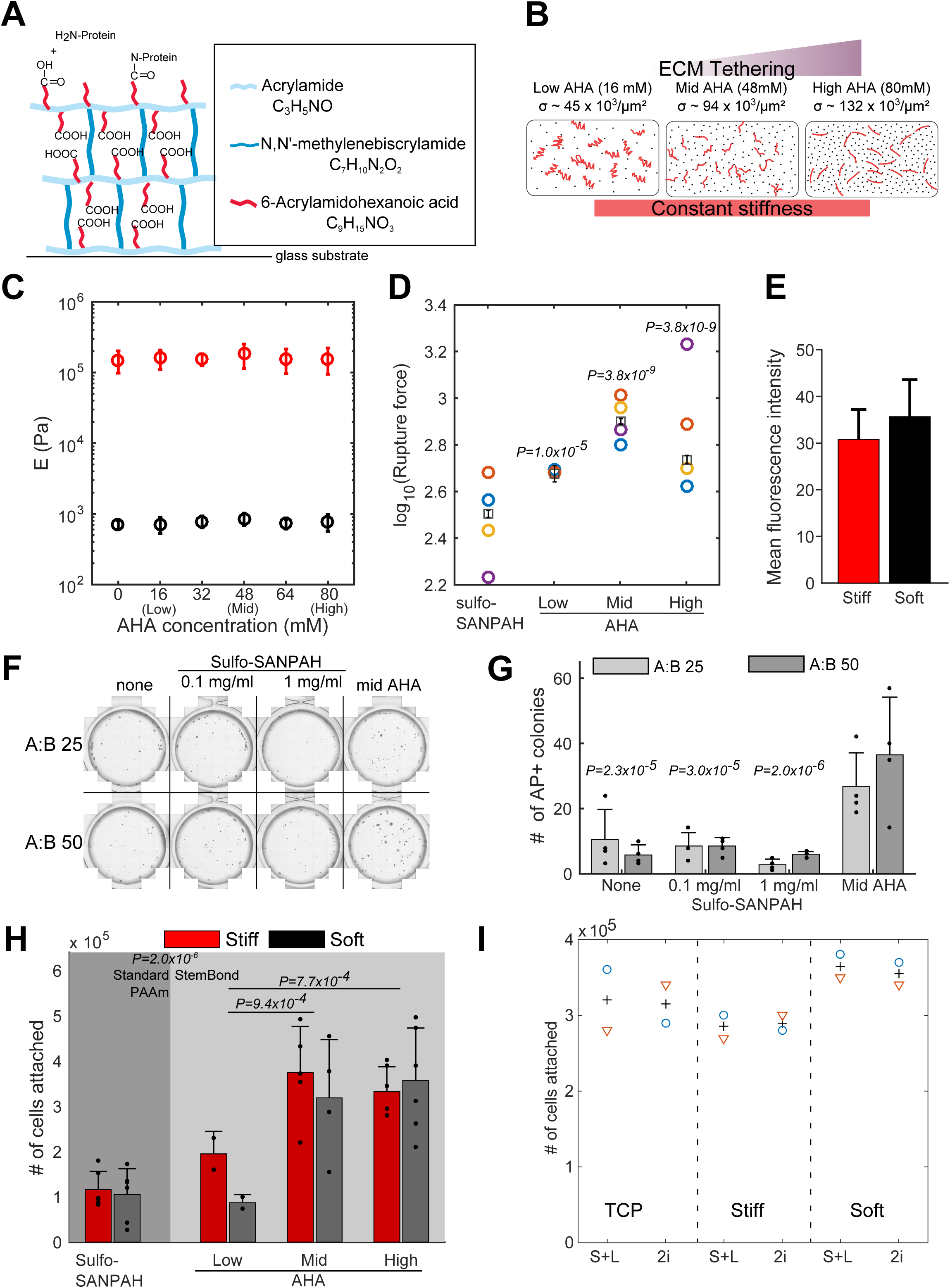
Novel gel surface chemistry & characterisation of StemBond hydrogels. A – StemBond hydrogels are bonded to a glass substrate by siloxane groups. The polyacrylamide hydrogel is synthesised with co-polymer 6-Acrylamidohexanoic acid (AHA). The AHA chains terminate with a carboxyl group which is used to bind primary amines of ECM proteins. B – Schematic of variations in strength of ECM tethering. Different AHA concentrations will give different surface densities σ of carboxyl groups (black dots) to which proteins can be covalently bonded. The estimated values of σ are given for 3 different AHA concentrations. Each ECM protein fibre (red lines) can create many covalent bonds so different AHA concentrations will lead to different strength in ECM tethering while not affecting the stiffness. C – Young’s Modulus of StemBond hydrogels with different proportions of acrylamide, bis-acrylamide and AHA co-polymer were measured using AFM nanoindentation. Young’s modulus in Pa, reported as mean over 5 replicates (30 indentations/gel) for soft (black) and stiff (red) hydrogels. D – Rupture force (log_10_) from significant binding events between fibronectin covalently bound onto different hydrogels and an AFM probe coated with anti-Fibronectin. Circles show average of individual experiments, squares and bars show mean +/− standard error from 2-way ANOVA linear model. P-values correspond to pairwise comparison to sulfo-SANPAH substrates. E – Quantification of protein coverage on StemBond hydrogels, estimated by coating gels with FITC-labelled BSA and measuring the mean fluorescence intensity at the gel surface (N ≥ 7). F – Snapshots of Alkaline-Phosphatase (AP) staining of ES colonies on soft hydrogels. Hydrogels were prepared with an acrylamide:bis-acrylamide ratio of either 25 (top row) or 50 (bottom row). Surface functionalisation is indicated by column headers. All hydrogels (including the unfunctionalised ones) were coated with 200µg/ml of fibronectin. Cells were seeded at clonal density for 6 days before fixation and staining. G– Mean counts of number of AP positive colonies from assay in D for different surface functionalisation methods (N = 4). P-values from ANOVA test comparing different functionalisation to Mid AHA hydrogels. A:B ratio was not a significant factor. H – ES cell attachment on standard PAAm hydrogels (left) and StemBond hydrogels (right). Standard PAAm hydrogels were functionalised using sulfo-SANPAH. Cells were counted after 48hrs in S+L (N≥2). P-values from (ANOVA test) comparing Low to Mid AHA, Low to High AHA and Sulfo –SANPAH gels to StemBond gel. (Note that all hydrogels were coated with the same concentration of ECM). Stiffness was not a significant factor (*p=0.58*). In all panels, unless otherwise noted, error bars represent standard deviation. I – Cell attachment on fibronectin-coated tissue culture plastic (TCP), stiff hydrogels and soft mid AHA hydrogels. 175’000 cells were plated for 24hrs in culture in 2i or S+L medium before being detached and counted (N = 2). ‘o’ and ‘Δ’ show individual data points, ‘+’ show mean.

We estimated that the surface density of ECM anchorage points approximately triples when AHA concentration is increased from 16mM (Low AHA) to 48mM (Mid AHA) and 80mM (high AHA) (**Figure 1B**). In order to measure ECM tethering strength, we used AFM cantilevers coated with an anti-fibronectin antibody. On both soft and stiff StemBond hydrogels, we detected significant binding events (**Figure S1A-C**). We did not, however, measure significant binding events on sulfo-SANPAH functionalised stiff hydrogels, primarily because the rupture length was very short on these gels (**Figure S1D**). The short rupture length on stiff sulfo-SANPAH substrates, which have relatively high ECM tethering density, likely indicates a weak tethering of ECM to the substrate. On soft sulfo-SANPAH substrates, the significant binding events had longer rupture lengths but significantly smaller rupture forces than on soft StemBond substrates (**Figure 1D and S1A-E**), also indicative of weaker ECM tethering to the substrate. Plateauing rupture forces on high AHA hydrogels suggest saturation of ECM tethering points. Notably, the rupture forces are very similar for both soft and stiff StemBond hydrogels (**Figure S1A and S1D**), and protein coverage levels were similar on soft and stiff substrates as confirmed by immunofluorescence (**Figure 1E**). Thus, in contrast to the widely used sulfo-SANPAH approach, controlling co-factor concentration on StemBond substrates allows strong, reproducible and tunable ECM tethering to the substrate, even on very soft PAAm hydrogels.

### StemBond hydrogels promote strong attachment of ES cells

We then tested whether stronger ECM tethering would improve ES cell adhesion to the hydrogels. We found, using two values of Acrylamide:Bis-acrylamide ratio (A:B) to vary pore size and two different concentrations of sulfo-SANPAH, that long-term cell attachment on fibronectin-coated StemBond hydrogels was significantly higher than on standard PAAm hydrogels functionalised with sulfo-SANPAH (**Figure 1F**). Notably, after a few days of culture on sulfo-SANPAH functionalised hydrogels, independent of concentration, there were clear floating colonies, and these hydrogels ultimately yielded as few attached colonies as the unfunctionalised hydrogels. In contrast, on StemBond hydrogels we observed many large colonies irrespective of A:B ratio (**Figure 1G**).

We then tested whether different levels of AHA would impact cell attachment over the course of 48 hours. We found that cell attachment was long-lasting on mid and high AHA hydrogels, and weaker on the low AHA hydrogels only (**Figure 1H**), which had similar attachment numbers to 1mg/ml sulfo-SANPAH hydrogels. Furthermore, substrate stiffness did not affect cell attachment on mid AHA and high AHA hydrogels, and cell attachment and viability were comparable to tissue culture plastic (TCP) (**Figure 1I and S1F**). Proliferation rates, as assessed by EdU incorporation, were approximately the same for all substrates (**Figure S1G**). Thus, StemBond hydrogels, in contrast to sulfo-SANPAH functionalised hydrogels, provide robust conditions for long-term culture of ES cells, even on soft substrates.

### Soft substrates enhance pluripotency

We first examined the consequences of substrate stiffness and ECM tethering on ES cells by performing RNA sequencing on ES cells cultured on StemBond hydrogels with three different ECM tethering densities and two substrate stiffnesses in S+L (**Table S1**). Morphologically, colonies on soft substrates were rounder, resembling naïve ES cells in 2i culture, whereas on stiff substrates colonies were flat and spread out, similar to ES cells on TCP (**Figure 2A**). ECM tethering density did not dramatically impact morphology, though we observed that colonies were systematically slightly less round on the high AHA soft substrates (**Figure 2A**). We also found that substrate stiffness had a greater influence on gene expression than ECM tethering density, with 452 differentially expressed genes due to changes in stiffness along with greater overall fold changes in gene expression, against only 52 differentially expressed genes due to ECM tethering (see **Figure 2B and Methods**). Principal component analysis confirmed that samples primarily cluster by substrate stiffness (**Figure 2C**). Gene ontology analysis of the differentially expressed genes revealed enrichment for stem cell population maintenance and transcription processes in genes upregulated on soft substrates, whereas cell adhesion and migration as well as some differentiation pathways were enriched in downregulated genes (**Figure S2A**). Immunostaining confirmed that markers of strong cell-ECM adhesion and of high cytoskeletal tension (mature, phospho-paxillin-rich focal adhesions and actin stress fibres) were diffuse or absent on soft substrates (**Figure S2B**).

**Figure 2:**
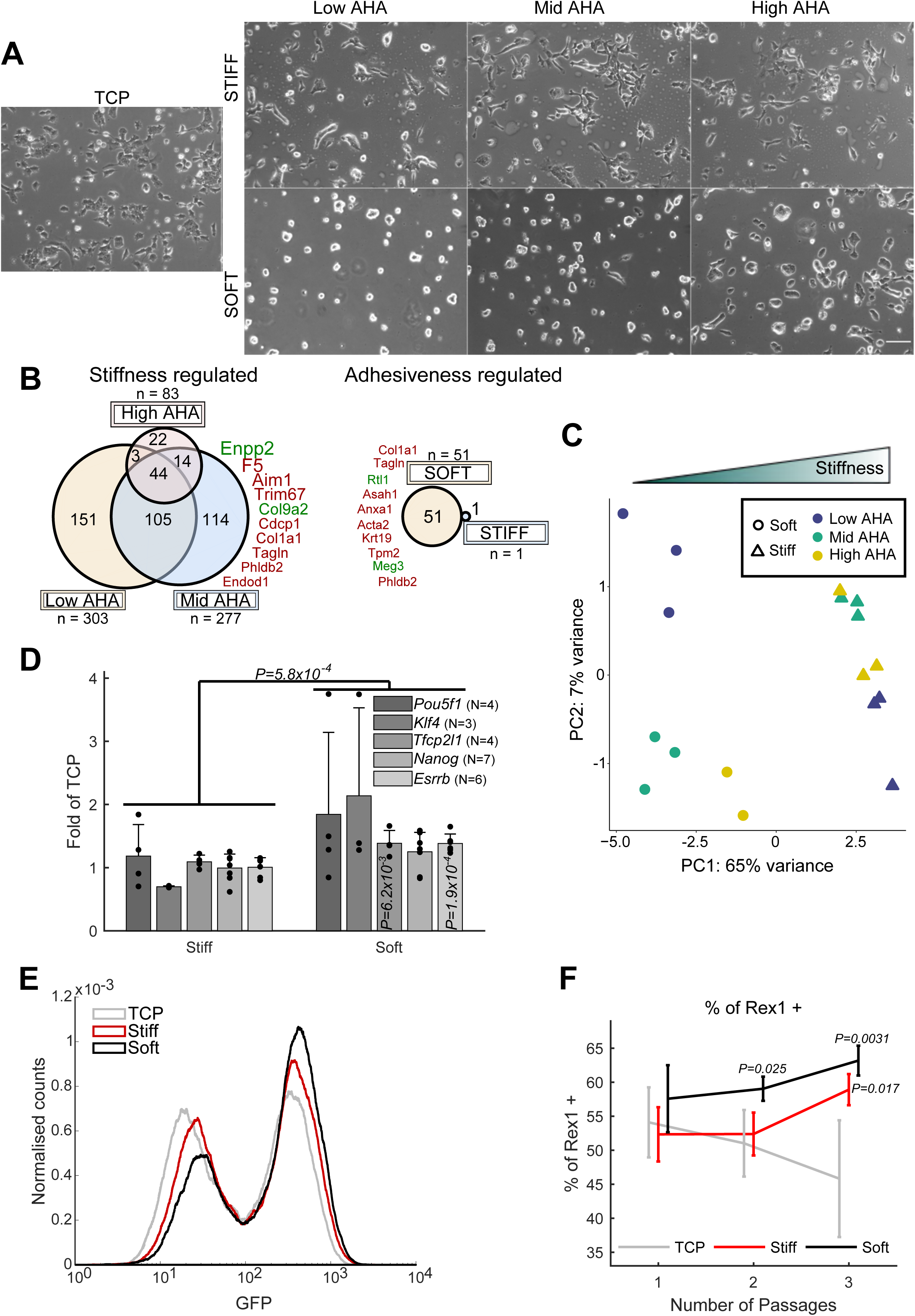
ES cell response to variations in substrate stiffness and ECM tethering. A – Brightfield images of cells after 24hrs in culture in S+L on tissue culture plastic (TCP), stiff (top row) and soft (bottom row) StemBond hydrogels with low AHA (left), mid AHA (centre) or high AHA (right). All surfaces coated with fibronectin. Scale bar: 100µm. B – Venn diagrams indicating the number of differentially expressed genes (padj < 0.05) in pair-wise comparisons. (Left) Number of genes regulated by substrate stiffness for low (n = 303), mid (n = 277) or high (n = 83) ECM tethering density. (Right) Genes regulated by ECM tethering on soft (51) and stiff (1) substrates. The top 10 genes with largest fold change are shown on the side of each diagram, with green/red indicating up-/down-regulation and font size proportional to the mean log_2_ fold change (soft compared to stiff for the stiffness-regulated genes or low/mid AHA compared to high AHA for the adhesiveness-regulated genes) C – Principal component analysis (PCA) plot computed on differentially expressed genes across all conditions (one point = one sample). Conditions are indicated by markers (stiffness) and colour-code (AHA concentration). D – Mean and standard deviation of mRNA expression of selected pluripotency genes on soft and stiff (mid AHA) substrates normalised to TCP. Cells were cultured on the substrates for 24hr in S+L. For each gene, p values indicate significant differences with TCP. The overall line and P-value shows there are significant differences due to stiffness across all targets tested (computed by n-way ANOVA). E – Flow cytometry profiles of reporter line Rex1GFP::d2 cultured on TCP (grey), stiff (red) or soft (black) hydrogels for 2 passages in S+L. For each condition, histograms of three replicate samples were averaged and smoothed. F – Average percentage of Rex1 positive cells determined from flow cytometry histograms (see **Figure 2E**). Cells were cultured in S+L on TCP (grey), and on soft (black) and stiff (red) high AHA substrates (N = 4). P-values are given where significant differences to TCP were found. In all panels error bars show standard deviation.

Using qPCR to assay for key naïve pluripotency transcription factors *Esrrb, Tfcp2l1, Nanog* and *Klf4*, we again found no significant differences between our low AHA substrates and high AHA substrates (**Figure S2C**). This finding confirms that ECM tethering density is not a significant factor in the regulation of pluripotency. In contrast, soft substrates led to an overall highly significant increase in the expression of *Esrrb, Tfcp2l1, Nanog* and *Klf4*, with, for example, 38% increase in the expression of *Tfcp2l1* and *Esrrb* in comparison to stiff substrates (**Figure 2D**). We conclude that while sufficient substrate adhesiveness is essential for the long-term attachment of ES cells, substrate stiffness is the key mechanical property impacting gene expression of ES cells.

There are two possibilities to explain the enrichment of naïve genes at the population level; either the population is more homogeneous, or naive genes are upregulated within each single cell. To test this, we used a destabilised GFP reporter line for *Rex1* (*Zfp42*), a high-fidelity reporter for naïve pluripotency (Kalkan et al., 2017), and measured fluorescence intensity over several days. We found that soft substrates significantly increased the proportion of *Rex1* positive cells but did not significantly shift the peak fluorescence of the naïve cells (**Figure 2E-F**). Additionally, this proportion increased over time in culture on both soft and stiff hydrogels (but not on fibronectin-coated plastic dishes) (**Figure 2F**). These data suggested that StemBond hydrogels – particularly the soft substrates – progressively improve the phenotypic homogeneity of ES cells in S+L conditions.

### Substrate stiffness induces genome-wide transcriptomic changes mirroring *in vivo* changes

To assess if the observed boost in naïve pluripotency would remain in conditions other than S+L, we performed RNA-sequencing in more challenging conditions, comparing gene expression of ES cells in serum with or without LIF and/or without the MEK/ERK inhibitor, PD03 (on mid AHA, soft and stiff substrates). We found that, in all media conditions, many genes were significantly differentially expressed between soft and stiff substrates, as indicated by the numbers in the Venn diagram (**Figure 3A**), with over 210 genes being modulated in more than 3 conditions. Regulation of stem cell pluripotency and PI3K pathways were two of the top enriched pathways identified by gene ontology analysis. The transcription of key mechanosensing structures, such as focal adhesions, cell-cell junctions, and the cytoskeleton, was also differentially regulated on soft and stiff substrates in all conditions (**Figure 3B and S3A-C**).

**Figure 3:**
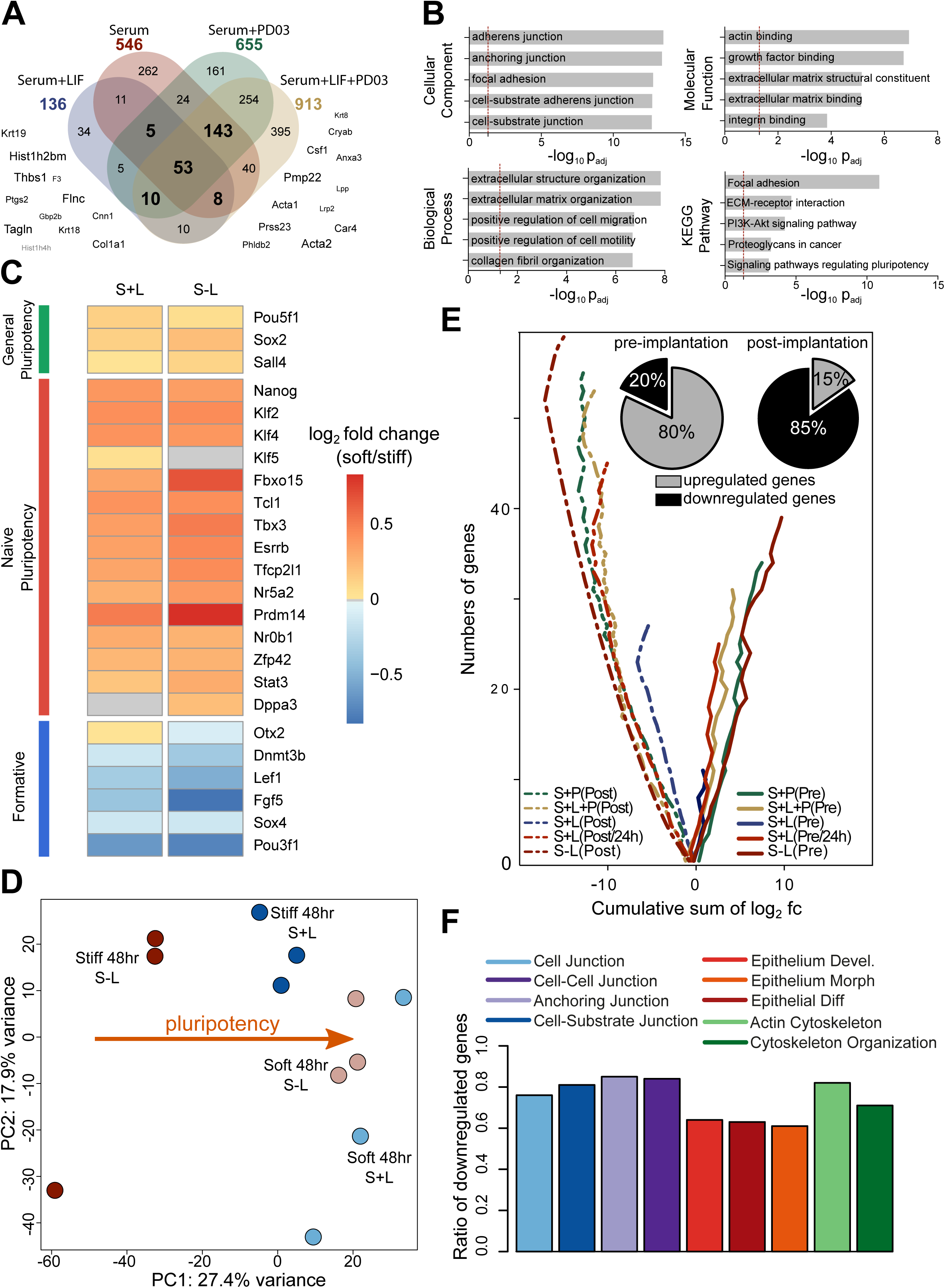
RNA sequencing shows stabilisation of pluripotency on soft substrates. A - Intersection of significantly modulated genes (p < 0.05, abs (log_2_fc)> 1, FPKM > 1) between soft and stiff substrates in all media conditions. In bold are the numbers of systematically modulated genes (*i.e.* in at least 3 conditions). Examples of some genes are given on the side with font size proportional to the average up/downregulation. log_2_fc = log_2_ fold change = log_2_(soft/stiff). B - Cellular components, biological processes, molecular functions and KEGG pathways that are enriched in the systematically modulated genes (n = 219). Dotted red lines represent the significant threshold (p < 0.05). See also **Figure S3**. C - Heatmap of log_2_fc for a selection of general pluripotency, naïve pluripotency and formative genes in different conditions. Conditions are S+L, 24hrs (‘S+L’) and serum without LIF, 48hrs (‘S-L‘). D - PCA plot for samples in S+L, 48hrs conditions based on the highly variable genes (n=2579, see also **Figure S5A**). The arrow indicates pluripotency potential with the more naïve pluripotent population clustered on the right hand side. E - Cumulative sum of log_2_fc for all genes (abs(log_2_fc) > 0.2) belonging to pre-implantation (solid lines) or post-implantation clusters (dotted lines). The diverging lines of the pre- and post-implantation genes indicate some systematic and opposing effect of substrate stiffness on those two clusters (if regulations were uncorrelated to substrate stiffness, solid and dotted lines would converge around zero cumulative log_2_fc). Pie charts show the average percentage of significantly up/downregulated genes over all conditions (percentages for each condition given in **Figure S5C**). S-L: serum without LIF; S+L: serum+LIF; L:LIF; P:PD03; Pre:pre-implantation; Post: post-implantation. F - Percentage of downregulated genes in serum without LIF condition for each of the selected biological processes. See also **Table S2**.

In detail, we found that the expression of naïve pluripotency factors, but not of general pluripotency factors, was significantly increased on soft compared to stiff hydrogels, whereas the expression of recently-defined formative genes (Kalkan et al., 2017; Smith, 2017), indicative of cells having exited naïve pluripotency, was significantly decreased on soft substrates (**Figure 3C and S4A-B and Table S2**). These changes in expression levels were observed across different media conditions (**Figure S4C-D**), and most pronounced in the absence of LIF and PD03. Notably, these changes were more pronounced after 24 hours in culture as opposed to 48 hours in culture, suggesting that phenotypic differences begin to be masked at high cell density. Principal component analysis (PCA) confirmed that cells on soft substrates cluster together independently of the presence or absence of LIF, while on stiff substrates cells cultured with and without LIF cluster separately (**Figure 3D and S5A**). These results were similar to differences observed in flow-cytometry and qPCR (**Figure 2E-F and S5B**), suggesting that soft substrates led to a LIF-independent expression of naïve pluripotency genes.

To understand if such differences in gene expression had biological significance, we compared them to the difference between pre- and post-implantation epiblasts, which have been characterised by sequencing early embryos (Boroviak et al., 2015). We computed the log_2_ fold change of the differentially expressed genes that were associated either to the pre- or post-implantation epiblast and found that among pre-implantation genes, 80% were upregulated on soft substrates, while among the post-implantation genes, 85% were downregulated on soft substrates (**Figure 3E and S5C, Table S2**). The trend was similar across all media conditions. Thus ES cells cultured on soft and stiff substrates recapitulated key transcriptional differences between pre-implantation epiblast and post-implantation epiblast *in vivo*, respectively.

The above results were most striking in serum without LIF (“S-L”), suggesting that ES cells on soft substrates are resistant to differentiation in serum conditions, even in the absence of LIF. Not only are formative genes lowly expressed, but pathways such as epithelialisation, essential to differentiation and, *in vivo*, to epiblast progression (Shahbazi et al., 2017), are impacted. We indeed found that, on average, 74% of differentially regulated components known to be involved in epithelialisation and cytoskeleton reorganisation were downregulated on soft substrates (**Figure 3F, Table S2**). These results further demonstrate that soft substrates provide a condition that is highly permissive to maintenance of naïve pluripotency.

### ES cell self-renewal in minimal media conditions

To test whether the observed transcriptional differences amounted to a functional gain in ES cells, *i.e.* an increased capacity for self-renewal, we performed clonogenicity assays. In these assays, we remove one or several self-renewing factor(s) for a defined period of time, then replate the cells at very low densities in 2i+LIF medium for 5 days to select for cells still able to self-renew in naïve conditions. Finally, we count the number of resulting naïve pluripotent colonies (**Figure 4A-i**). First, we removed LIF (a positive effector of self-renewal) from the medium for 5 days on TCP, stiff and soft StemBond substrates. On TCP, removal of LIF led to drastic loss of self-renewal. On stiff substrates, there was a modest loss of naïve pluripotency upon removal of LIF. On soft substrates, however, significantly more cells maintained naïve pluripotency than on stiff substrates and TCP (**Figure 4B**). Surprisingly, twice as many cells maintained naïve pluripotency on soft substrates without LIF when compared to the control of undifferentiated cells grown on TCP in S+L. These results suggest that in serum-only conditions, StemBond hydrogels at least partially compensate for LIF to sustain self-renewal of ES cells.

**Figure 4:**
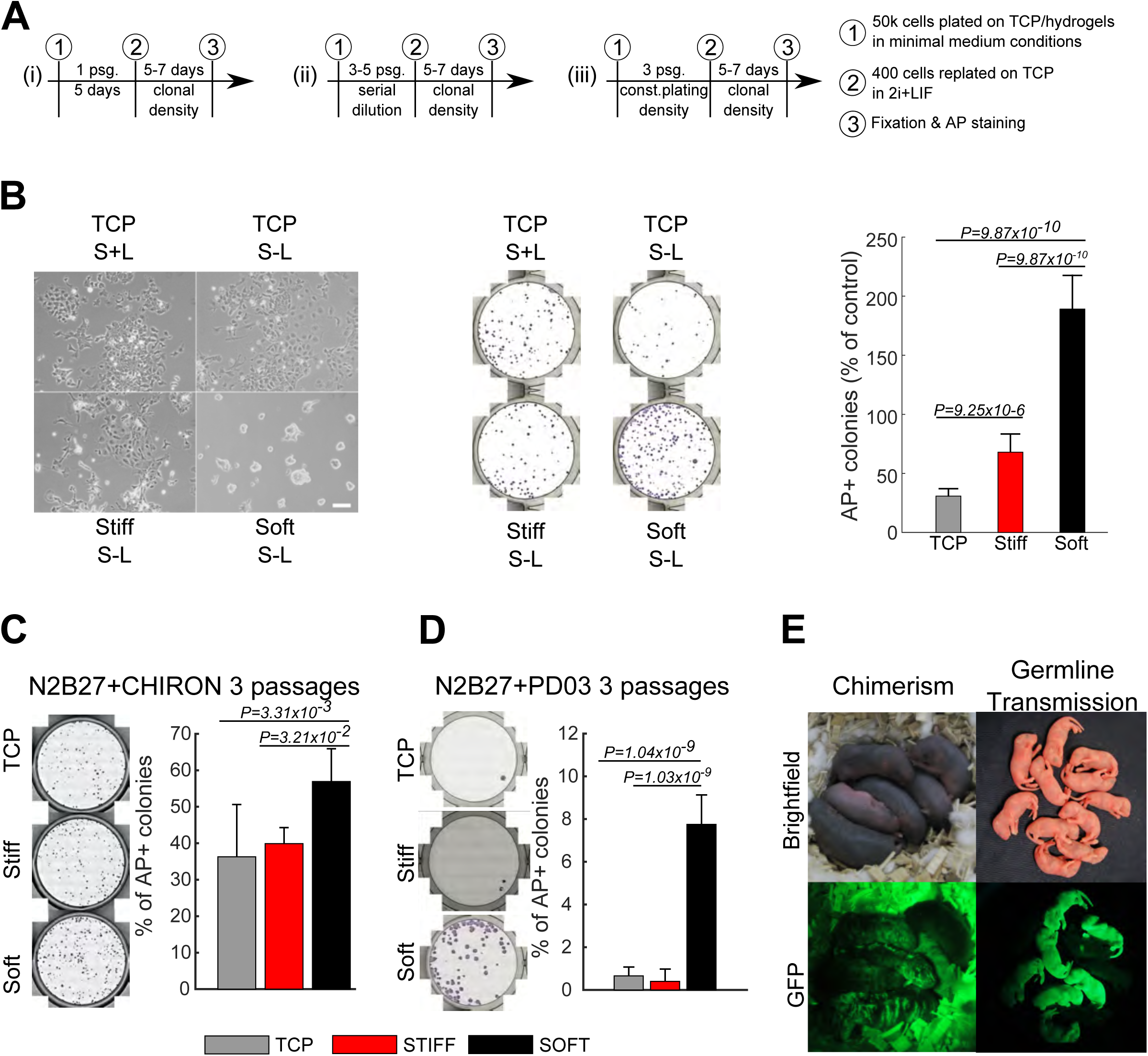
Clonogenicity assays show that soft gels enhance self-renewal in minimal conditions. A - Procedure for clonogenicity assays in minimal medium. First, cells were seeded on different substrates, in varying medium conditions. Cells were either (i) left for 5 days in S-L or (ii) & (iii) passaged every 2-3 days. Passaging (psg) was done either (ii) by serial dilution from 1:3 to 1:4 or (iii) by keeping a constant plating density of 50,000 cells per substrate. Second, cells were replated at clonal density (100 cells/cm^2^) on gelatin-coated plastic in 2i+LIF to allow self-renewing cells to form clonal colonies. Third, colonies were fixed and stained for alkaline phosphatase (AP). B – Clonogenicity assay of cells after 5 days in S-L on different substrates following protocol (i). (Left) Brightfield images showing the morphology of cells after 5 days in different conditions. Scale Bar: 100µm. (Centre) Snapshot of the stainings for AP. (Right) Quantification of the number of AP positive colonies in % of the number retrieved in control (undifferentiated) conditions (S+L, on TCP substrates) (N=8). C – Clonogenicity assay after cells were cultured for 3 passages in N2B27+CHIRON on different substrates following protocol (iii). (Left) Snapshots of stainings for AP. (Right) Quantification of the number of AP positive colonies in % of number of plated cells (N ≥ 5). D – Clonogenicity assay after cells were cultured for 3 passages in N2B27+PD03 on different substrates following protocol (iii). Cells here were from a C57BL/6-Agouti background. (Left) Snapshots of staining for AP. (Right) Corresponding quantification of the number of AP positive colonies in % of number of plated cells (N=8). E – First (left) and second (right) generation chimeric mice obtained after the injection into blastocysts of GFP-transfected cells cultured on soft substrates in N2B27+PD03 for 3 passages following protocol (ii). Note that the high background signal in the chimerism GFP image is due to autofluorescence of the bedding, but clear GFP+ regions are visible on mice.

We then tested whether soft substrates could compensate for a single of the two otherwise essential self-renewing inhibitors in 2i, CHIRON (GSK3β inhibitor) or PD03 (MEK inhibitor), thus allowing propagation of ES cells for multiple passages in minimal media conditions. For this, we cultured cells on different substrates for 5 passages in N2B27+CHIRON by serial dilution, before replating them at very low density in 2i+LIF to assess how many cells were naïve pluripotent (**Figure 4A-ii**). Significantly more cells gave rise to naïve colonies from soft substrates than from stiff substrates (**Figure S6A-B**). Moreover, cell survival was also significantly higher on soft substrates (**Figure S6C**). Therefore, since the higher survival could be a confounding factor because it influences plating density in a serial dilution experiment, we then repeated the experiment keeping constant cell plating density at each passage (**Figure 4A-iii**). After 3 passages of constant plating density, there was no cell survival on stiff substrates, but survival was still high on soft substrates. Furthermore, we found in these conditions that there were significantly more naïve pluripotent cells on soft substrates than on stiff substrates and TCP (**Figure 4C).**

In N2B27+PD03, cell survival was very low on all substrates, so the serial dilution clonogenicity assay was not possible. Therefore, we performed the assay with constant cell plating density over multiple passages (**Figure 4A-iii**). This allowed propagation for up to 3 passages on soft and stiff substrates only, with no cells surviving past passage 2 on TCP (**Figure S6D**), in line with previous studies (Wray et al., 2010). Survival did gradually decrease on soft and stiff substrates, with no significant differences in cell numbers, and less than 10% of cells giving rise to naïve colonies on both substrates (**Figure S6E-F**). Because sensitivity to MEK/ERK inhibition (but not to GSK3β inhibition) has been shown to depend on cell line (Wray et al., 2010), we repeated this assay with a different cell line, from a C57BL/6-Agouti background. In this case, we found a greater effect of the substrate stiffness on the number of naïve pluripotent colonies obtained with ~9% on soft substrates but only ~1% on stiff substrates and TCP (**Figure 4D**). We conclude that soft substrates more robustly provide a microenvironment supportive of self-renewal in N2B27+PD03.

To ultimately confirm that the cells cultured in N2B27+PD03 on soft hydrogels were still naïve pluripotent, we injected GFP-targeted ES cells into mouse blastocysts to test their ability to contribute to the germ layers and the germline. We obtained 5 viable chimaeras from the first injection and 7 GFP-expressing pups from the next generation, demonstrating germline transmission (**Figure 4E**). To our knowledge, this is the longest that ES cells have ever been successfully maintained in this highly challenging condition. Taken all together, we conclude that soft StemBond hydrogels improve ES cell self-renewal with only one of the 2 otherwise essential inhibitors, CHIRON or PD03.

Given that soft substrates are highly conducive to the maintenance of naïve pluripotency even in the presence of minimal chemical support, we next cultured ES cells in basal medium, N2B27 only, for 3 days. In these conditions, unlike the others, there is no chemical support for maintenance of naïve pluripotency, therefore ES cells normally begin to differentiate. Surprisingly, we found that there were very few naïve colonies returned from the clonogenicity assay for all substrate conditions. Furthermore, we used the Rex1::GFPd2 cell line to follow cell differentiation in N2B27, and found that *Rex1* was downregulated within 48 hours with similar dynamics on all three substrates (**Figure S6G**). We conclude that in pure differentiation conditions, exit from naïve pluripotency proceeds normally regardless of substrate condition. This finding suggests that soft substrates do not block differentiation, but instead synergise with other naïve pluripotency signals such as PD03, CHIRON, serum and LIF to boost the maintenance of ES cell self-renewal.

### Soft StemBond hydrogels promote the acquisition of naïve pluripotency

Bespoke microenvironments such as 3-dimensional matrices have been shown to accelerate cell reprogramming, in part through increased epigenetic remodelling (Caiazzo et al., 2016). We thus hypothesised that soft StemBond hydrogels could not only promote the maintenance, but also enhance the induction of naïve pluripotency. We started from primed epiblast stem cells (EpiSC), which can be reprogrammed to naïve induced pluripotent stem cells (iPSC) by the forced expression of a single naïve transcription factor together with signalling cues (Brons et al., 2007; Guo et al., 2009; Tesar et al., 2007). We employed doxycycline-inducible *Esrrb* (iEsrrb) as an efficient driver of reprogramming in EpiSCs (Festuccia et al., 2012; Stuart et al., 2019, 2014) with a Rex1::dGFP-IRES-bsd reporter for the naïve identity (Kalkan et al., 2017). EpiSCs did not stably adhere to sulfo-SANPAH treated substrates and only weakly to mid AHA StemBond hydrogels, but they adhered well to high AHA StemBond hydrogels, which we thus used for reprogramming assays. We plated EpiSCs on plastic, stiff and soft substrates for 24 hours before inducing reprogramming with 2i+doxycycline and then assessed the efficiency of the process at different time points (**Figure 5A**). After 24 hours, cells were more clustered together on soft substrates than on TCP and stiff hydrogels (**Figure 5B**). Both early markers of reprogramming, *Tfcp2l1* & *Klf2* and the late marker *Zfp42* (Rex1) showed a twofold increase in gene expression on soft substrates. The general pluripotency marker *Pou5f1* (Oct4), which transiently drops when reprogramming is triggered, was more effectively maintained on soft substrates (**Figure 5C**). These results suggested a boost in the naïve pluripotency network during the reprogramming process. Correspondingly, we found a significantly higher end-point efficiency of reprogramming on soft substrates, with twice as many naïve colonies on soft substrates at day 8 (**Figure 5D**). Naïve genes were expressed at similar levels in resultant iPSCs demonstrating full reprogramming (**Figure S7**). Therefore, soft StemBond hydrogels not only support ES cell self-renewal but also promote specification of naïve pluripotency during reprogramming.

**Figure 5:**
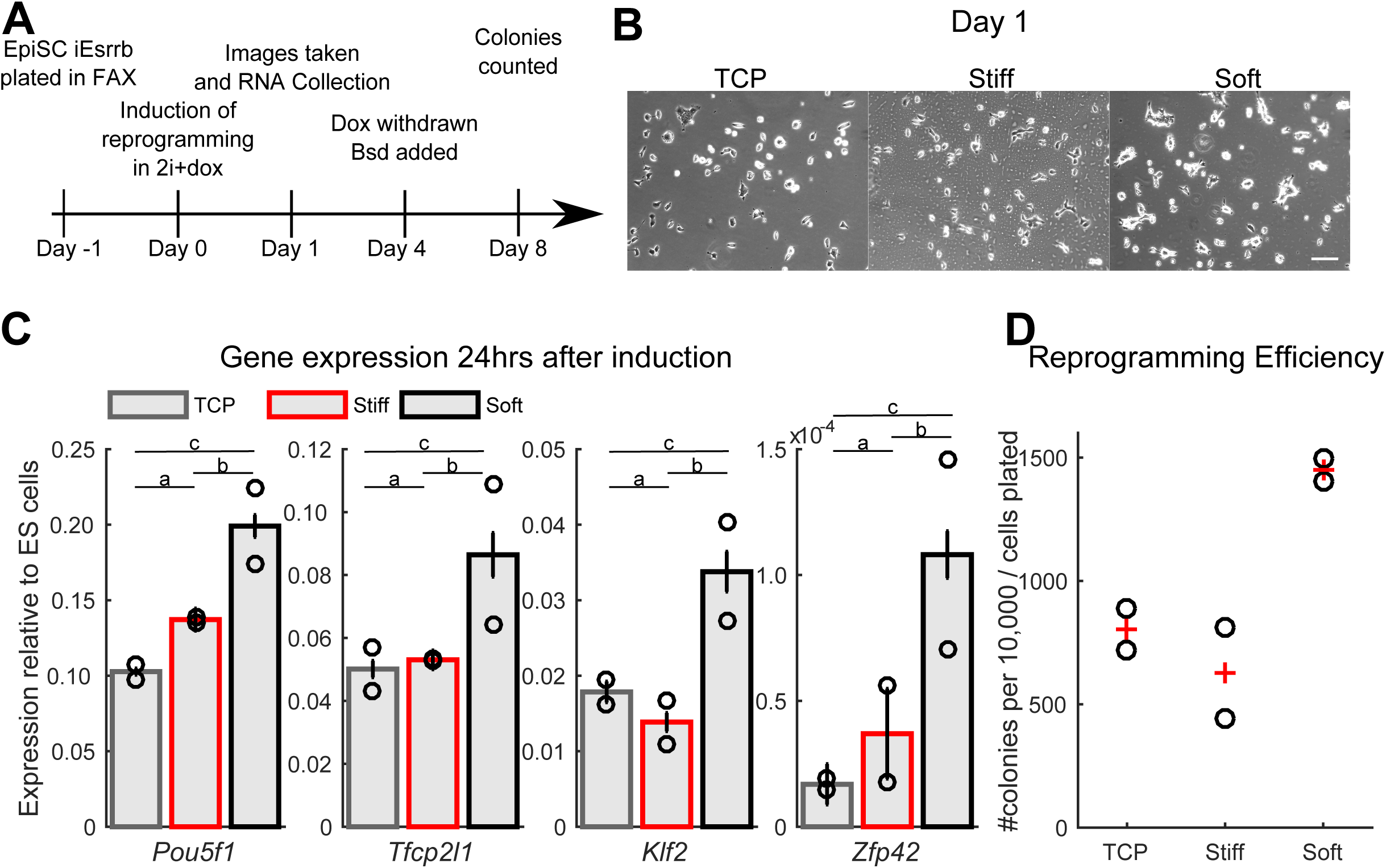
EpiSC reprogramming boosted on soft substrates. A - Schematic of reprogramming procedure. EpiSCs were plated in their maintenance Fgf2+ActivinA+XAV (FAX) medium. *iEsrrb*-driven reprogramming was induced by switching to 2i + doxycycline (dox) the next day. After 4 days of reprogramming, dox was withdrawn and blasticidin (bsd) was added to select for naive colonies with an active *Rex1* promoter. Blasticidin-resistant iPSC colonies were counted on day 8. B – Phase contrast images (magnification 10x) 24 h after induction of *iEsrrb* EpiSC reprogramming in 2i+dox on TCP, stiff and soft substrates. Scale bar, 100µm. C – Gene expression profile 24hrs after induction of *iEsrrb* EpiSC reprogramming in 2i+dox. Expression is presented relative to *Gapdh* then normalised to 2iLIF naïve ES cells levels. Circles represent individual data points, bars show average (N = 2). a,b,c, indicate groups that were compared in a non-parametric Friedman test over all genes in panel: a: *P* = 0.9243, b: *P* = 0.0942; c: *P* = 0.0373. D – Number of iPSC colonies counted on day 8 on TCP, stiff and soft substrates, as a measure for reprogramming efficiency, circles show individual data points, ‘+’ show average (N = 2). See also **Figure S7**. In all panels error bars show standard deviation.

### Substrate stiffness acts on both LIF/JAK/STAT3 and FGF/MEK/ERK pathways in ES cells

To gain insights into how soft substrates enhanced the maintenance and acquisition of naïve pluripotency, we investigated core signalling pathways known to regulate the naïve state. In particular, we investigated MEK/ERK signalling, which triggers exit from naïve pluripotency and supports lineage commitment (Kunath et al., 2007; Nichols & Smith, 2009), as well as LIF/JAK/STAT3 signalling, which supports naïve self-renewal (Niwa et al., 1998). We found that the signalling activity of these pathways depended on substrate mechanics, with significantly lower ERK activity and higher STAT3 activity on soft substrates compared to stiff substrates (**Figures 6A-B**), suggesting that soft substrates stabilise naive pluripotency both by boosting self-renewal through enhancing STAT3 activity and by reducing ERK-driven differentiation.

**Figure 6:**
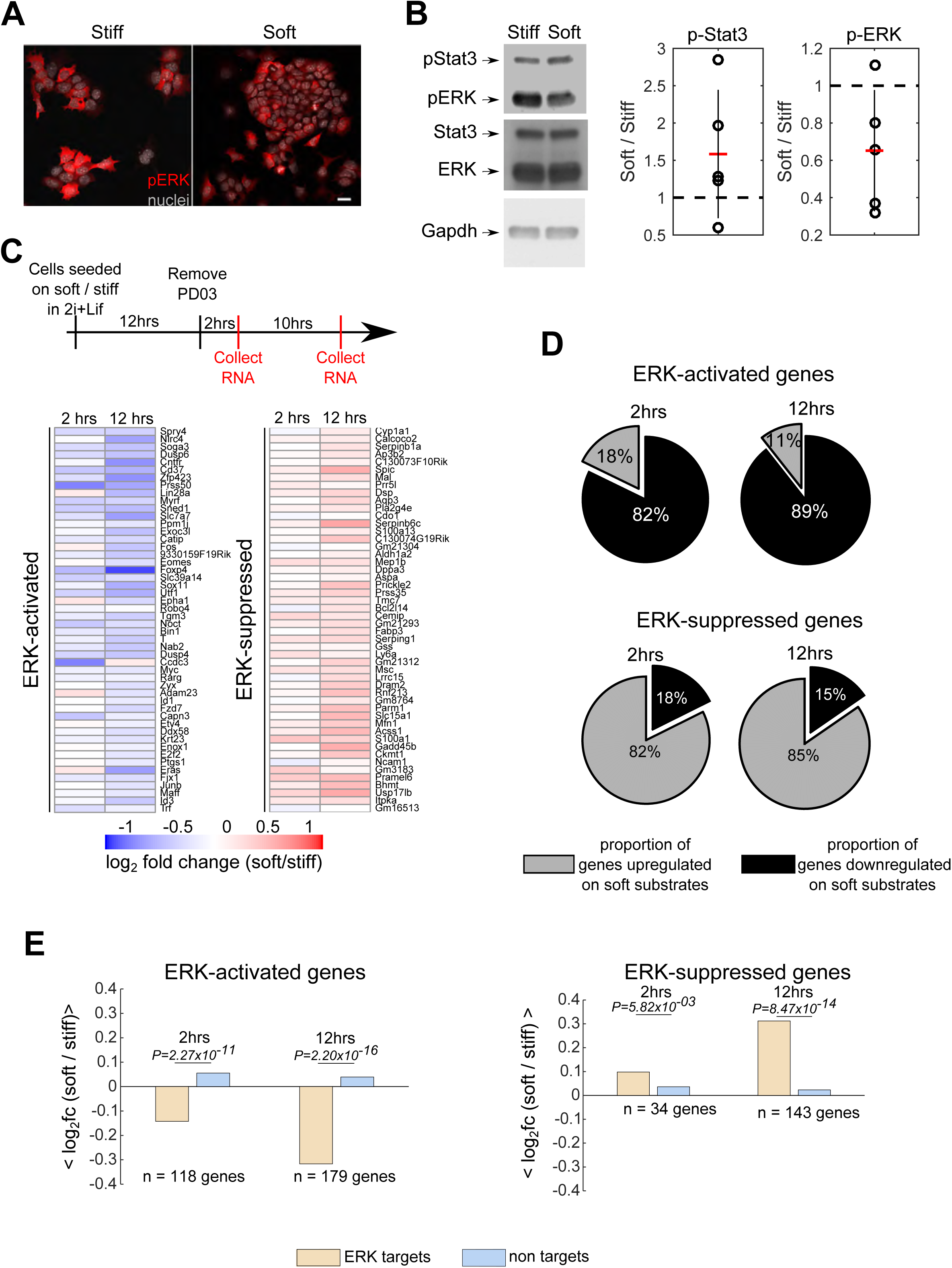
MEK activity is substrate stiffness dependent. A – Immunofluorescence for phospho-ERK (red) and DAPI (grey) on stiff (left) and soft (right) substrates. Cells were seeded on the gels for 24hrs in S+L. Scale bar 20µm. B – (Left) Western blots for phospho-STAT3, phospho-ERK, total STAT3 and total ERK from cells grown on soft and stiff gels for 24hrs in S+L. GAPDH was used as loading control. (Right) Mean +/− std of the ratio of intensity of soft/stiff after normalisation to GAPDH (N = 5). C – (Top) Design of experiments for RNA-sequencing indicating the timing of the different steps and PD03 removal. Experiments performed in triplicate. (Bottom) Heatmaps of log_2_ fold change in expression between soft and stiff substrates after PD03 removal. (Left) Top 50 activated genes and (right) Top 50 suppressed genes. The genes listed are those which are co-regulated after inhibitor removal on both soft and stiff and show the largest fold change on stiff substrates when compared to control in 2i+LIF. D - Pie charts giving the percentage of genes whose expression was higher on soft (grey) or on stiff (black) at 2hrs (left) and 12hrs (right) after removing PD03. Percentages are relative to all the ERK-regulated genes at 2hrs, and to the ERK-regulated genes with|log_2_ (t=12hrs/t=0hr)| > 1 at 12hrs. (E) Average log_2_ fold change of expression between soft and stiff for the ERK-regulated genes, compared to the same number of a random selection of non-regulated genes. P-values from Wilcox rank sum are given above each comparison.

ERK regulates gene expression both by phosphorylating cytoplasmic substrates and by binding DNA motifs of target genes (Yang, Sharrocks, & Whitmarsh, 2013). To test whether substrate mechanics changed ERK‘s transcriptional activity (Paszek et al., 2005; Trappmann et al., 2012) we then looked for mechanosensitive transcriptional targets of ERK. To do this, we first carried out RNA-sequencing at early time-points (2hrs and 12hrs) after removing PD03 from 2i+LIF media (**Figure 6C and Table S3**). Note that CHIRON+LIF media is sufficient to maintain naïve pluripotency, so this assay tests specifically for the effects of ERK signalling induction in ES cells independent of changes in pluripotent state. Known ERK targets such as Dual-Serine Phosphatases (*Dusps*) and Immediate-Early Genes (*Fos, Egr1, Jun*) were all activated after 2 hours on soft and stiff substrates, confirming that PD03 withdrawal led to ERK activation in all conditions.

Moreover, we observed a systematic effect of substrate stiffness on the magnitude of ERK-target regulation (**Figure 6C**). A comparison of the expression of all co-regulated genes (*i.e.* either activated or suppressed by ERK on both substrates) revealed that nearly 90% of ERK-activated genes were downregulated on soft substrates, whereas 85% of ERK-suppressed genes were upregulated on soft substrates compared to stiff ones (**Figure 6D**). This was already evident 2 hours after removing PD03, and significantly more so at 12 hours. The fold change was significantly greater for these ERK-regulated genes than for a random selection of non-regulated genes (**Figure 6E**). As a control, we performed the same analysis removing CHIRON from 2i+LIF. Both canonical and non-canonical Wnt pathway genes are suppressed on soft and stiff substrates as CHIRON is removed, with no dependence on substrate-stiffness in those GSK3β-mediated changes, even after 12 hours (**Figure S8 and Table S4**). Taken together, these results suggest that substrate stiffness significantly impacts the activation and transcriptional activity of the ERK pathway, which provides a mechanism for the optimised maintenance of self-renewal on soft substrates.

## DISCUSSION

In this study, we established ES cell culture with StemBond hydrogels, which are two-dimensional substrates that can straightforwardly replace TCP for stem cell culture. The challenge here was to have sufficient substrate adhesiveness – independent of substrate stiffness – to ensure robust cell-ECM adhesion on one hand, and both self-renewal and differentiation on the other hand. The most commonly used method of PAAm hydrogel functionalisation – sulfo-SANPAH – did not allow for stable ES cell attachment even at high concentrations. Interestingly, poor cell attachment was observed on both soft and stiff substrates, indicating that it is the instability of sulfo-SANPAH substrate crosslinking, rather than pore size (Trappmann et al., 2012; Wen et al., 2014), that is responsible for poor cell attachment. Our results thus call into question the suitability of culturing ES cells on sulfo-SANPAH functionalised hydrogels, and other stem cells, particularly from soft tissue, should be studied in this context.

By co-polymerising the functional group together with acrylamide, allowing a controlled, robust and adjustable ECM tethering density on soft and stiff substrates, StemBond hydrogels overcome the limitations presented by sulfo-SANPAH functionalisation. We found that intermediate and high concentrations of AHA led to high ECM tethering strength, mirroring results of adhesion assays which also found that those conditions were more stable for culture of ES cells than low AHA or sulfo-SANPAH hydrogels. In the future, other methods of PAAm functionalisation should be tested to see how tethering strength is affected.

Previous studies on different types of substrates suggested that intermediate levels of cell-fibronectin interactions (Hunt, Singh, & Schwarzbauer, 2012), low cell-ECM traction forces (Chowdhury, Li, et al., 2010; Hayashi et al., 2007) and limited cell spreading (Murray et al., 2013) promote self-renewal. Our results are in line with these, showing that weak cell-ECM adhesion does not enhance self-renewal, whereas substrates with low stiffness - even with high ECM tethering - do enhance self-renewal. Indeed, substrate stiffness was found to be a potent regulator of the naïve identity. On soft hydrogels, the expression of naïve pluripotency genes and of genes associated with the pre-implantation epiblast was upregulated, independently of substrate adhesiveness, both with and without LIF. These transcriptional changes reflected a more homogeneous population, and were coupled with an increase in the ES cells’ self-renewal capacity in the absence of LIF. Furthermore, the doubling of the reprogramming efficiency of EpiSC in 2i without LIF argues for the fact that soft substrates are not only permissive for the maintenance, but also for the acquisition, of naïve pluripotency.

This permissivity is most likely due to synergies between soft substrates and exogenous signalling cues (serum, LIF, CHIRON or PD03). We indeed found that in the absence of any pluripotency-driving signal, ES cells will differentiate on all substrates. On TCP or stiff substrates, two of these signals are required to ensure self-renewal of naïve cells, because one of these signals alone would lead to differentiation or cell death. In contrast, we here show that soft substrates in conjunction with a single pluripotency-driving signal (serum, PD03 or CHIRON) promote maintenance and acquisition of naïve pluripotency. Taken together, these results suggest that ‘softness’ constitutes an axis for maintaining naïve pluripotency, and acts in coordination with chemical cues to steer stem cell fate. Future work should elucidate the molecular nature of the interplay between soft substrates and signalling pathways.

Overall, we conclude that by promoting strong attachment but low cytoskeletal tension, soft StemBond hydrogels provide an optimal microenvironment for long term culture of ES cells and EpiSC, and are well suited for controlled studies of interplay between mechanical signalling and differentiation in stem cells. This work opens new possibilities to define alternative culture conditions with better control over stem cell fate decisions in physiologically-mimetic conditions and with minimal need of small molecule inhibitors. Importantly, StemBond hydrogels can be adapted to match mechanical and adhesive properties other stem cell niches, as has been done recently to reverse the ageing of rat oligodendrocyte progenitor cells (Segel et al., 2019). Future work will expand the scope of StemBond substrates to multiple stem cell types and culture systems.

## ACKNOWLEDGEMENTS

We are grateful to Peter Humphreys for assistance with imaging and imaging analysis, Sally Lees and staff for tissue culture work, Maike Paramor and staff for preparation of sequencing libraries, Michael A. Barber for alignment of sequencing data, G. Chu and staff for animal husbandry, Carla Mulas for comments on the manuscript, Ivan B. Dimov and Amelia Joy Thompson for help with AFM measurements. Rosa26-CreERT2+/+ cells were a kind gift from the lab of Bon-Kyoung Koo. Illumina RNA sequencing was performed at the CIGC, Cambridge.

This work was financially supported by core funding from the Medical Research Council and Wellcome to the Wellcome-MRC Cambridge Stem Cell Institute, and individual grants to KJC and JCRS from the Medical Research Council (MR/M011089/1) and to KJC from the European Research Council (772798). BXT is supported by a Wellcome four-year PhD grant. JCRS is a Wellcome Trust Senior Research Fellow (WT101861). KJC is a Royal Society University Research Fellow. KF received support by MRC Career Development Award (G1100312/1) and ERC (772426).

## MATERIALS AND METHODS

### Cell culture

E14 mouse Embryonic stem cells and Rex1GFPd2 cells, a kind gift from Austin Smith’s laboratory at University of Cambridge, were cultured in either 10% Foetal Calf Serum + LIF medium (S+L) or 2i+LIF medium following established protocols. S+L was supplemented with 2mM L-glutamine (Gibco), MEM non-essential amino acids (Gibco), 1mM Sodium Pyruvate (Invitrogen), and 0.1mM 2-mercaptoethanol (Gibco). 2i+LIF was made up of N2B27 defined basal medium (1:1 Neurobasal and DMEM/F-12 medium (Invitrogen), 0.5% N2 (homemade), 1% B27 (ThermoFisher Scientific), 2mM L-glutamine (Gibco), 0.1mM 2-mercaptoethanol (Gibco) supplemented with MEK inhibitor (1 µM PD0325901), GSK3β inhibitor (3 µM CHIR99021) and/or 0.2µg/ml murine LIF as indicated. Rex1+/*dGFP-IRES-bsd TetOn-Esrrb + CAG-rtTA3* (*iEsrrb* (Stuart et al., 2019)) EpiSCs were cultured in N2B27 as above, supplemented with 12.5 ng/ml Fgf2, 20ng/ml ActivinA (Hyvonen lab, Cambridge) and 6.25 µg/ml XAV 939 (Tocris). EpiSCs and reprogramming experiments were conducted in hypoxic conditions (7% CO2 and 5% O2).

ESCs, iPSCs and EpiSCs were dissociated with accutase (Millipore) during passaging. Cells were plated on either tissue culture plastic (TCP) or hydrogel substrates coated with 200µg/ml human plasma fibronectin (Corning, NY, USA and Millipore, Germany) at a density of 5,000 to 15,000 cells/cm^2^.

### Staining and imaging

Cell fixation, staining and slide mounting were done according to (Agley, Rowlerson, Velloso, Lazarus, & Harridge, 2013) (see Table 3 for antibody list). Samples were imaged on Leica TCS SP5 confocal microscope or Zeiss LSM 710 confocal microscope. Images were then analysed using Leica software and ImageJ. To assess gel surface coating, we used FITC-labelled BSA (Sigma-Aldrich).

### Flow cytometry

The GFP signal in Rex1GFPd2 cells was monitored by flow cytometry on a Dako Cytomation CyAn ADP high-performance unit. Results were analysed using FlowJo and custom scripts in Matlab (Mathworks).

In Figure 2E-F, cells were plated on high AHA StemBond hydrogels in S+L and passaged every 48hrs (keeping the same density at each passage).

### Hydrogel fabrication

Prior to hydrogel fabrication, coverslips were cleaned and functionalised with either Bind-Silane (GE Healthcare) for support coverslips or 20% Surfasil in chloroform (Fischer Scientific) for top coverslips. Hydrogel solutions were prepared according to Table 1 and polymerised between a support and a top coverslip for 15min. The density of ligand binding sites was tuned by adapting the concentration of the co-polymer 6-acrylamidohexanoic acid (AHA) (IUPAC name 2-(prop-2-enoylamino)hexanoic acid). Table 1 gives recipes of soft and stiff hydrogels for a final concentration of 48mM AHA and A:B ratio of 25, which was used throughout the study, unless otherwise noted. After polymerisation, the top coverslips were removed and gels were rinsed twice in methanol, and soaked in PBS.

**Table 1:** Hydrogel recipes for 48mM co-factor concentration. 2M AHA stock solution was prepared in methanol.

| E (kPa) | Acrylamide 40% | Bis-acrylamide 2% | AHA 2M | H<sub>2</sub>O | TEMED | APS 10% |
| --- | --- | --- | --- | --- | --- | --- |
| 0.75 | 35 µl | 30 µl | 12 µl | 415.5 µl | 2.5 µl | 5 µl |
| 160 | 200 µl | 150 µl | 12 µl | 130.5 µl | 2.5 µl | 5 µl |

Hydrogels were then activated by 30min treatment in 0.2M EDAC (Sigma-Aldrich, Germany) + 0.5M NHS (Sigma-Aldrich, Germany) in MES buffer, pH 6.1. They were coated with 200µg/ml fibronectin diluted in HEPES buffer (50mM pH 8.5) overnight at 4°C. After washes, gels were blocked with 0.5M Ethanolamine in HEPES buffer. Gels were stored at 4°C until use.

Hydrogels without AHA were activated with sulfo-SANPAH (ThermoScientific, MA,USA). 1mg/ml (unless otherwise noted) Sulfo-SANPAH was dissolved in HEPES, and activated by UV-light for up to 30min.

### Atomic Force Microscopy

#### Substrate’s stiffness

Atomic Force Microscopy (AFM) was used to measure the elastic modulus of StemBond hydrogels, on a Nanowizard CellHesion 200 AFM (JPK Instruments) placed on an inverted microscope (Axio Observer A1, Zeiss) fitted with a motorised stage. Polystyrene beads (microParticles, Germany) with a diameter of 37.28µm (for soft hydrogels) and 10.28µm (for stiff hydrogels) were glued onto tipless silicon cantilevers (Arrow-TL1, Nanoworld, Switzerland) using MBond 610 Adhesive (Micro-Measurements). Spring constants of 0.03 – 0.07 N/m for soft hydrogels, 0.1 – 0.3 N/m for stiff hydrogels were determined by the thermal noise method. 10 force-distance curves at 3 different locations per gel (x5 replicate gels) were measured. Post-processing and analysis were done using JPK SPM data processing software, using the Hertz model to fit the approaching curve and extract the values of the substrate’s Young’s modulus.

#### Matrix tethering strength

Si-N gold-coated cantilevers with pyrex-nitride pyramidal tip (Nanoworld, #PNP-DB) were coated following protocol of (Chirasatitsin & Engler, 2010; Wen et al., 2014). Briefly probes were cleaned for 30s in chloroform, then incubated with 5M ethanolamine-HCl overnight. They were then washed in PBS and incubated for 30min in 25mM BS3 (bis(sulfosuccinimidyl)suberate, ThermoFisherScientific, #21580), washed again in PBS and immersed in 200µg/ml antibody solution for 30min. Finally probes were rinsed and kept at 4°C until use. Antibodies used were (rb) anti-Fibronectin (ABCAM, ab2413) and (rb) anti-IgG (Cell Signalling, #2729S). Gels were prepared either with co-factor AHA or activated with sulfo-SANPAH. A control had no activation, and no coating. Activated gels were coated overnight with fibronectin at 200µg/ml unless specified otherwise, then extensively rinsed. All gels were passivated with 1% BSA. Gels were stored in PBS at 4°C until use. 2 gels per condition were probed for both batches of measurements. Each sample was probed over 3 regions of 10×10 grids, points spaced by 10 to 20 µm. Head speed was 5µm/sec, loading rate ~ 160nN/s setpoint was at 500pN. Dwell time at the surface was 1-3sec before retraction. Data was processed using the JPK Data Processing Software (Version 6.1.131) and Matlab (Mathworks). Briefly, for each measurement, we plotted the rupture force (minimal value of the vertical deflection) against the rupture length (see **Figure S1A**). To rule out non-specific interactions, we used probes coated with anti-IgG. Data on stiff and soft substrates were analysed separately because of differences in sample/probe interaction area (setpoint was kept constant at 500 pN). Processing then followed these steps:

i. For each curve, rupture force and rupture length are extracted
ii. A threshold of both rupture force and rupture length were set at the median of the distributions for negative controls.
iii. The 2-d density map of negative controls and of samples were computed (**Figure S1B**). The regions where the density of samples was more than twice that of samples were defined as clusters of significant events, after a median filtering to smooth the cluster (**Figure S1C**).
iv. For all the significant binding events, the mean rupture force is determined (see **Figure 1D**), and the overall fraction of significant binding events (compared to all measurements done on each samples in **Figure S1E**). Graphs show average over 2-4 samples, errors are standard error of the mean. Total number of measurements per sample ranged from 34 to 696 and number of significant events per sample ranged from 0 to 233.

### Gene expression assays

For gene expression analysis, cells were lysed in RLT buffer (QIAGEN) and RNA extracted using RNEasy extraction kit (Qiagen). RT-PCR was performed using Superscript Transcriptase III (ThermoScientific, MA, USA) and gene expression was then assessed using the relevant Taqman probes (FAM) (Table 2) and Gapdh (VIC) as endogenous control.

**Table 2:** List of probes for gene expression assays

| Gene | Cat No | Company |
| --- | --- | --- |
| Esrrb | Mm00442411_m1 | ThermoFisher Scientific |
| Gapdh (Endogenous Control) | 4352339E | ThermoFisher Scientific |
| Nanog | Mm02384862_g1 | ThermoFisher Scientific |
| Lefty2 | Mm00774547_m1 | ThermoFisher Scientific |
| Klf4 | Mm00516104_m1 | ThermoFisher Scientific |
| Klf2 | Mm00500486_g1 | ThermoFisher Scientific |
| Nr0b1 | Mm00431729_m1 | ThermoFisher Scientific |
| Pou5f1 | Mm00658129_gH | ThermoFisher Scientific |
| Tfcp2l1 | Mm00470119_m1 | ThermoFisher Scientific |
| Zfp42 | Mm03053975_g1 | ThermoFisher Scientific |

### RNA Sequencing and Analysis

Library preparation was done by in-house facility using Pico mammalian V2 (Takara), NuGen (NuGen, CA,USA) or RiboZero and Nextflex (Bioo Scientific, TX, USA) kits. Sequencing was performed on Illumina HiSeq4000 yielding 350 Million reads per lane.

### RNA sequencing data processing, transcriptome analysis and network analysis

#### Dataset of Figure 2, S2

Mouse genome build GRCm38/mm10 were used to align reads with GSNAP version 2015-09-29 (Wu & Nacu, 2010). Genes were annotated using Ensembl release 81 (Cunningham et al., 2015) and read counts were quantified using HTSeq (Simon Anders, Pyl, & Huber, 2015). Differential expression analysis was computed using DESeq2 (Love, Huber, & Anders, 2014) on protein coding genes, from pairwise comparisons of samples with either same AHA concentration or same stiffness.

DAVID 6.7 (Huang, Sherman, & Lempicki, 2009b, 2009a) was used to compute the statistical enrichment of Gene Ontology terms, using genes up-and down-regulated by stiffness as input. Cytoscape (Shannon et al., 2003) and the enrichment map plugin (Isserlin, Merico, Voisin, & Bader, 2014) were used for network construction and visualisation (**Figure S2A**). Node size is scaled by the number of genes contributing to over-representation of biological processes; edges are plotted in width proportional to the overlap between gene sets. Node colour represents the different AHA conditions in which each biological process is significantly enriched.

#### Dataset of Figures 3, S3 – S5

Mouse genome build GRCm38/mm10 were used to align reads with STAR 2.5.2a (Dobin et al., 2013), 2013). Genes were annotated using mouse annotation from Ensembl release 87 (Yates et al., 2016) and splice junction donor/acceptor overlap settings were tailored to the read length of each dataset. Alignments to gene loci were quantified with Htseq-count (Simon Anders et al., 2015) based on annotation from Ensembl 87.

Principal component analyses was performed based on log_2_ FPKM values computed with the Bioconductor packages DESeq (S Anders & Huber, 2010), FactoMineR (Lê, Josse, & Husson, 2008) in addition to custom scripts. In addition DESeq was used to perform differential analysis between stiff and soft substrates.

In order to identify genes with the greatest expression variability we fitted a non-linear regression curve between average log_2_ FPKM and the square of coefficient of variation. Specific thresholds were applied along the x-axis (average log_2_ FPKM) and y-axis (log CV^2^) to identify the most variable genes.

DAVID 6.8 (Huang et al., 2009b, 2009a) was used to compute the statistical enrichment of Gene Ontology terms, using modulated genes in Serum-only conditions as input (**Figure 3F**).

STRING database (Snel, Lehmann, Bork, & Huynen, 2000) was used to retrieve gene-gene interaction and Cytoscape was used to visualise the resulting network (**Figure S3B**). Transcription factor and transcription co-factor annotation were downloaded from AnimalTFDB (http://bioinfo.life.hust.edu.cn/AnimalTFDB/).

Cytoscape (Shannon et al., 2003) and the enrichment map plugin (Isserlin et al., 2014) were used for network construction and visualisation (**Figure S3C**). Node size is scaled by the number of genes contributing to over-representation of biological processes; edges are plotted in width proportional to the overlap between gene sets. The color represents the centered percentage of up/downregulated genes for each biological process.

#### Dataset of Figure 6

Mouse genome build GRCm38/mm10 were used to align reads with Tophat v2.1.0 (Kim et al., 2013). Genes were annotated using Ensembl release 86 (Yates et al., 2016) and read counts were quantified using Featurecount v1.5.0 (Liao, Smyth, & Shi, 2014). Differential expression analysis was computed using DESeq2 (Love et al., 2014) on protein coding genes.

### Clonogenicity Assays and Alkaline Phosphatase staining

For clonogenicity assays in S+L, 50,000 cells were plated on the hydrogels in the indicated medium for 5 days (**Figure 4A-I** and **4B**). For clonogenicity assays following multiple passaging in N2B27+CHIRON and N2B27+PD03, cells were cultured on the hydrogels for 3 passages by serial dilution or keeping constant plating density as indicated in **Figure 4A** and in figure legends. Then 400 cells/well (**Figure 4C**) or 1000 cells/well (**Figure 4D**) were replated on gelatin-coated TCP in 2i+LIF for a minimum of 5 days. Cells were subsequently fixed in 8% formaldehyde and stained using the Alkaline-Phosphatase kit (86R 1KT, Sigma-Aldrich, Germany) following the manufacturer’s protocol. 4x images were acquired using CellSens software and an X-51 Olympus microscope system with motorised stage and camera. Colonies were then segmented and counted using ImageJ with manual verification at each step. In some wells there were colonies lining the side of the coverslip but beneath the hydrogel and these were therefore not counted.

### Chimeras

This research has been regulated under the Animals (Scientific Procedures) Act 1986 Amendment Regulations 2012 following ethical review by the University of Cambridge Animal Welfare and Ethical Review Body (AWERB). Use of animals in this project was approved by the ethical review committee for the University of Cambridge, and relevant Home Office licenses (Project license No. 80/2597) are in place.

Rosa26-CreERT2+/+ mouse embryonic stem cells from a C57BL/6-Agouti background (gift from Koo lab) were first transfected with a piggyback transposon vector (PB-GFP). Cells were then cultured in N2B27+PD03 on soft hydrogels for 3 passages by serial dilution of 1:2 at each passage. The cells were then dissociated into single cells, before injection into C57BL/6 host blastocysts at stage E3.5. Contribution of the injected cells to the mice is reflected by GFP expression. 5 out of 6 pups from the first generation of chimeras had significant GFP expression. 7 pups from the second generation expressed GFP (**Figure 4E**), demonstrating germline contribution.

### Reprogramming of EpiSCs

10,000 EpiSCs per well (6-well plate) were plated in N2B27+Fgf2+ActivinA+XAV medium, on the fibronectin-coated TCP and hydrogels. 24 hours later, reprogramming was induced by medium switch to N2B27+2i+ 1 µg/ml doxycycline (MP Biomedicals). After 4 days, dox-induction of *iEsrrb* was withdrawn and 20μg/ml blasticidin (Gibco) was applied to select for *Rex1*::dGFP-IRES-bsd reporter activity. On day 8, 4x images were acquired using CellSens software and an X-51 Olympus microscope system with motorised stage and camera. iPSC colonies with active *Rex1* reporter were counted manually. Resultant iPSCs were passaged once onto TCP then RNA lysates were harvested.

### Western Blots

Cells were lysed in RIPA buffer (Cell Signaling, Leiden, The Netherlands) and spun down. The supernatant was then denatured in SDS buffer at 95°C for 5min. Gradient mini-protean gels 8%-14% (Bio-Rad) were used for the western blots. Protein gels were then transferred onto a nitrocellulose membrane using a standard wet transfer procedure. Membranes were subsequently blocked with 5% BSA in TBS-Tween 20 for 2hrs, before overnight staining with primary antibodies at 4°C. Secondary antibodies incubated for 1hr at room temperature. ECL Fire (ThermoScientific, MA, USA) and ECL Prime (GE Healthcare) reagents were used to reveal the blots. See table 3 for antibody list.

**Table 3:** List of antibodies

| Antibody | Cat No | Company | Lot number | Dilution |
| --- | --- | --- | --- | --- |
| phospho ERK | 4370 S | Cell Signaling | #17 Ref:05/2016<br>#17 Ref:01/2017 | 1:1000 (WB), 1:200 (IF) |
| ERK | 4695 S | Cell Signaling | #21 Ref:05/2016 | 1:1000 (WB) |
| phospho Stat3 (m) | 41135 S | Cell Signaling | #5 Ref:07/2016 | 1:1000 (WB) |
| phospho Stat3 (rb) | 91455 | Cell Signaling | # 22 Ref:12/2013 | 1:1000 (WB) |
| Stat3 | 9139 S | Cell Signaling | #10 Ref:12/2016 | 1:1000 (WB) |
| Gapdh (m) | 97166 S | Cell Signaling | #3 Ref:05/2017 | 1:2000 (WB) |
| Gapdh (rb) | 5174 S | Cell Signaling | #6 Ref:11/2016 | 1:2000 (WB) |
| anti rb HRP | 7074 S | Cell Signaling | #26 Ref:12/2016 | 1:2500 (WB) |
| anti m HRP | 7076 S | Cell Signaling | #32 Ref:12/2015 | 1:2500 (WB) |
| phospho Paxillin | 2541 | Cell Signaling | #6 Ref:08/2015 | 1 :50 (IF) |
| Alexa-Phalloidin<br>555 | 8953 | Cell Signaling | #3 Ref:11/2016 | 1 :20 (IF) |

### Statistical Tests

In all figures, error bars show standard deviation (std) over N independent replicates (N indicated in legend), unless otherwise indicated.

Statistical differences across substrate conditions were determined using the ANOVA test on independent samples (tests done using Matlab). In most figures, tests are designed to compare across different types of substrate (TCP, Stiff and Soft hydrogels); in Figures 1D & 1G, multiple adhesiveness levels are included in the design. In **Figure 2D**, an n-way ANOVA was used to test for the significance of stiffness over multiple genes. Results are indicated in the figures with the convention throughout that *, **, *** indicate p < 0.05, p < 0.01 and p < 0.001 (no ‘*’ or n.s. means no statistically significant differences were found). In **Figure 5C**, the non-parametric Friedman test was used to compare substrate effects across the panel of 4 genes.

For **Figure 6E**, a non-parametric test was used (Wilcoxon rank sum, wilcox.test in R) over n genes, to determine if the expression of a selection of genes was systematically different on stiff versus soft substrates, compared to a random selection of the same number of genes.

### Estimation of surface density of adhesive ligand anchoring points

The surface density σ of anchoring points was estimated by the following formula: σ = (*N_A_⋅* [*AHA*])^2/3^ ⋅ 10^−10^ (Nsites/µm^2^) where N_A_ is the Avogadro constant, and [AHA] is the molar concentration of AHA.

## SUPPLEMENTARY DATA

### Table S1: Results of RNA sequencing, dataset of Figure 2

Normalised counts per gene for all samples that passed quality control. Outlier samples with high % of ERCC spike-in reads were discarded. Genes with less than 2 counts across all samples were discarded. Normalised counts (counts per million mapped counts) were obtained using the fpm function of DESeq2, with the robust option, i.e. using the size factors to normalise. Rows are labelled with EnsemblID, columns are labelled by sample stiffness. The samples are named by stiffness (“Soft” or “Stiff”) and adhesiveness (“16” = LowAHA, “48” = MidAHA and “80” = HighAHA).

### Table S2: Results of RNA sequencing, dataset of Figure 3

<u>Sheet “Table”</u> lists the genes differentially regulated between soft and stiff substrates in different medium conditions. For each comparison, the four columns indicate mean expression (FPKM) on soft, mean expression (FPKM) on stiff, log2 fold change (soft/stiff) and adjusted p-value.

<u>Sheet “Clusters”</u> lists the genes belonging to the pre- and post-implantation clusters used for Figure 3E.

<u>Sheet “Enrichment Score”</u> lists the pathways linked to post-implantation analyzed in Figure 3F (in Serum only conditions). For each pathway, the columns indicate the p-value for the pathway being significantly enriched, the enrichment score, the number of genes differentially expressed (k), the number of genes in the pathway annotation (K), the relative number of differentially expressed genes (k/K), and the ratio of number of genes downregulated on soft substrates over upregulated on soft substrates.

### Table S3: Results of RNA sequencing, dataset of Figure 6

Normalised counts for all protein coding genes for all samples during removal of PD03 (see also **Figure 6**). Normalised counts (counts per million mapped counts) were obtained using the fpm function of DESeq2, with the robust option, i.e. using the size factors to normalise. Rows are labelled per Gene Name. Genes with no counts across all samples were discarded. Columns are labelled per sample name which indicates the substrate (“Soft”/“Stiff”), the time after PD03 removal (“t0”,“t12”=12hrs,“t2”=2hrs) and the replicate number (1 or 2).

### Table S4: Results of RNA sequencing, dataset of Figure S8

Normalised counts for all protein coding genes for all samples during removal of CHIRON. One sample with too low number of reads was discarded. Normalised counts (counts per million mapped counts) were obtained using the fpm function of DESeq2, with the robust option, i.e. using the size factors to normalise. Rows are labelled per Gene Name. Genes with no counts across all samples were discarded. Columns are labelled per sample name which indicates the substrate (“Soft”/“Stiff”), the time after CHIRON removal (“t0”,“t12”=12hrs,“t2”=2hrs) and the replicate number (1 or 2).

**Figure S1:**
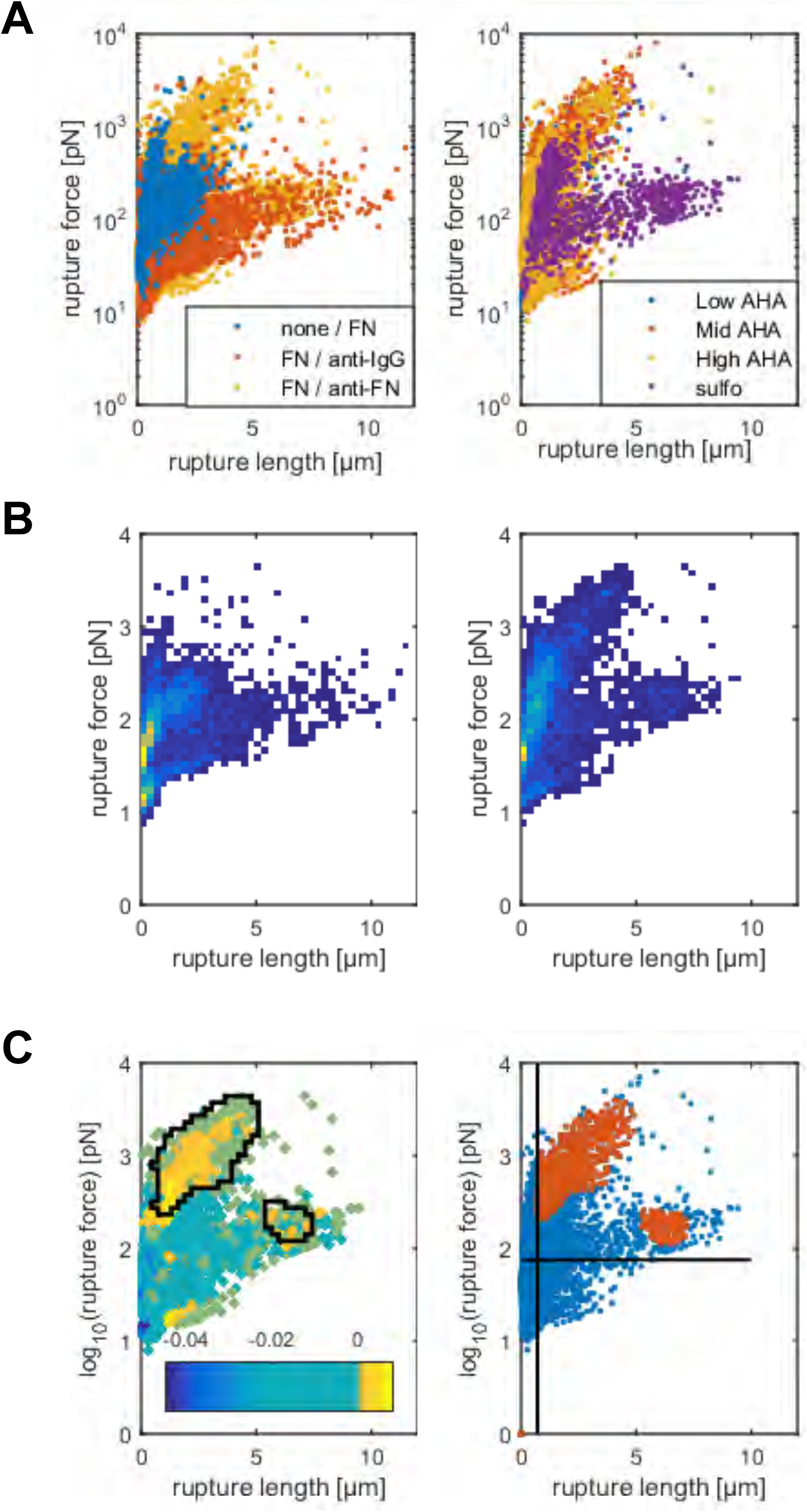
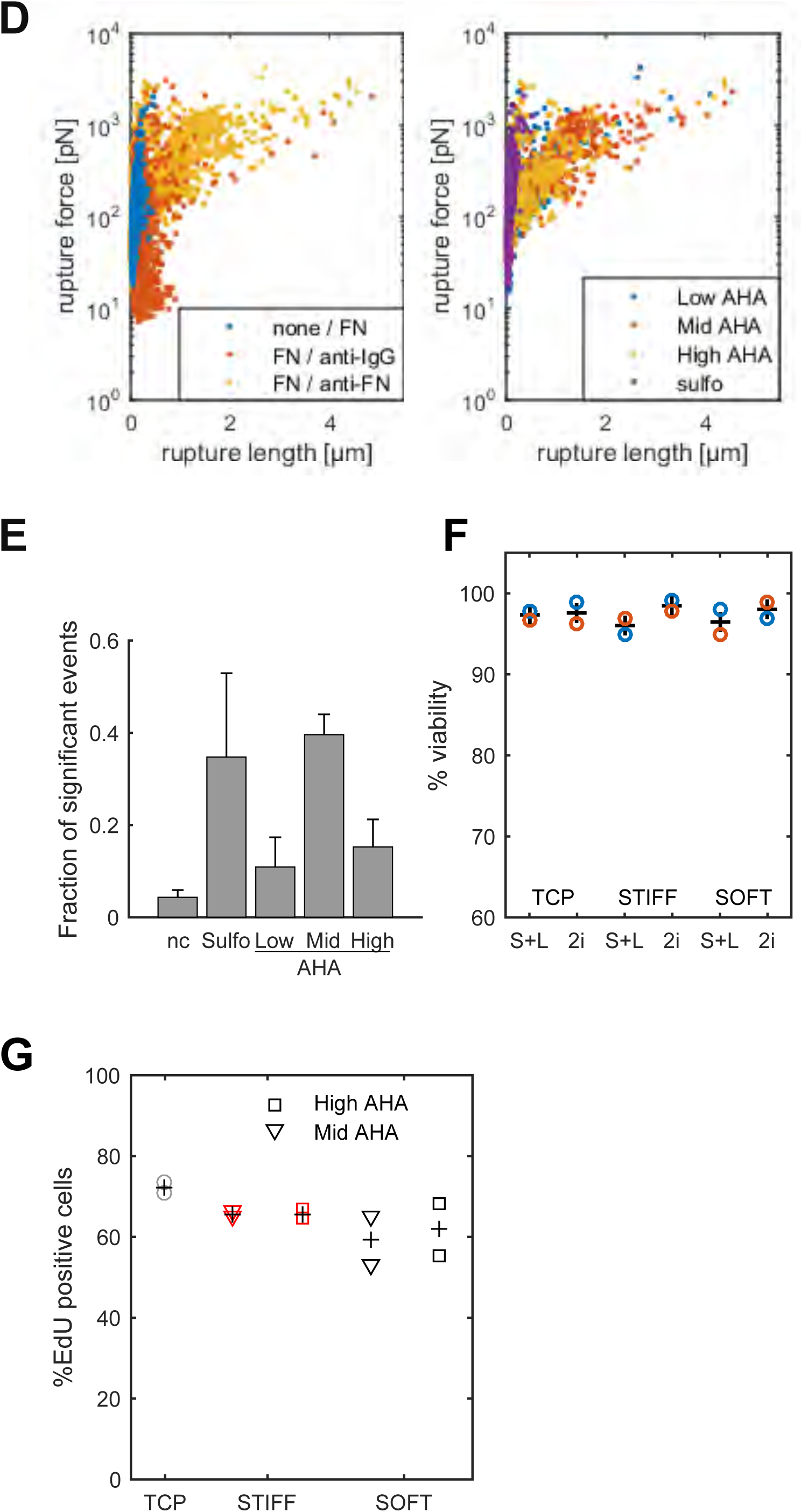
in relation to Figure 1. A – Plots showing rupture force vs rupture length for all measured curves on soft hydrogels. Each point corresponds to one measurement. (Left) Comparison of measurements for substrates coated with either fibronectin (‘FN’) or no matrix protein (‘none’) and probed with a cantilever functionalised either with anti-Fibronectin (‘anti-FN’) or anti-IgG antibody. Anti-IgG antibody serves to detect non-specific binding events. (Right) Comparison of measurements for different substrate functionalisation between 1mg/ml sulfo-SANPAH or Low, Mid and High AHA StemBond hydrogels. B – Density maps of the all measurements for negative controls (left) and real samples (right). C – Same data as in A (right panel). (Left)The colour code shows the amplitude of density (samples)-2xdensity(controls). This was used to identify clusters of points where sample measurements did not overlap with negative controls. The clusters of significant events are outlined in black (see **Methods** for details). (Right) Similar to the left panel, red points are within the clusters of significant events, blue points are out and discarded for following steps. Vertical and Horizontal lines show the median of rupture length and rupture force of negative controls, used as thresholds. D – Plots showing rupture force vs rupture length for all measured curves on stiff hydrogels. Each point corresponds to one measurement. (Left) Comparison of measurements for substrates coated with either fibronectin (‘FN’) or no matrix protein (‘none’) and probed with a cantilever functionalised either with anti-Fibronectin (‘anti-FN’) or anti-IgG antibody. Anti-IgG antibody serves to detect non-specific binding events. (Right) Comparison of measurements for different substrate functionalisation between 1mg/ml sulfo-SANPAH or Low, Mid and High AHA StemBond hydrogels. E– Fraction of significant binding events for each type of substrate functionalisation on soft hydrogels. nc: negative controls, Sulfo: 1mg/ml sulfo-SANPAH. Bars show mean +/− standard error over 2-4 replicate samples. F - Percentage (individual measurements ‘o’ and mean ‘+’) of viable cells assessed by Trypan Blue staining counted on flow cytometer (ViCell) after 24 hrs on different substrates (N = 2). G – Percentage of Edu+ cells. Cells were seeded overnight in 2i on gels, then incubated for 25min with EdU before fixing and quantification by imaging. ‘Δ’ and ‘○’ show individual measurements, ‘+’ show the mean (N = 2).

**Figure S2:**
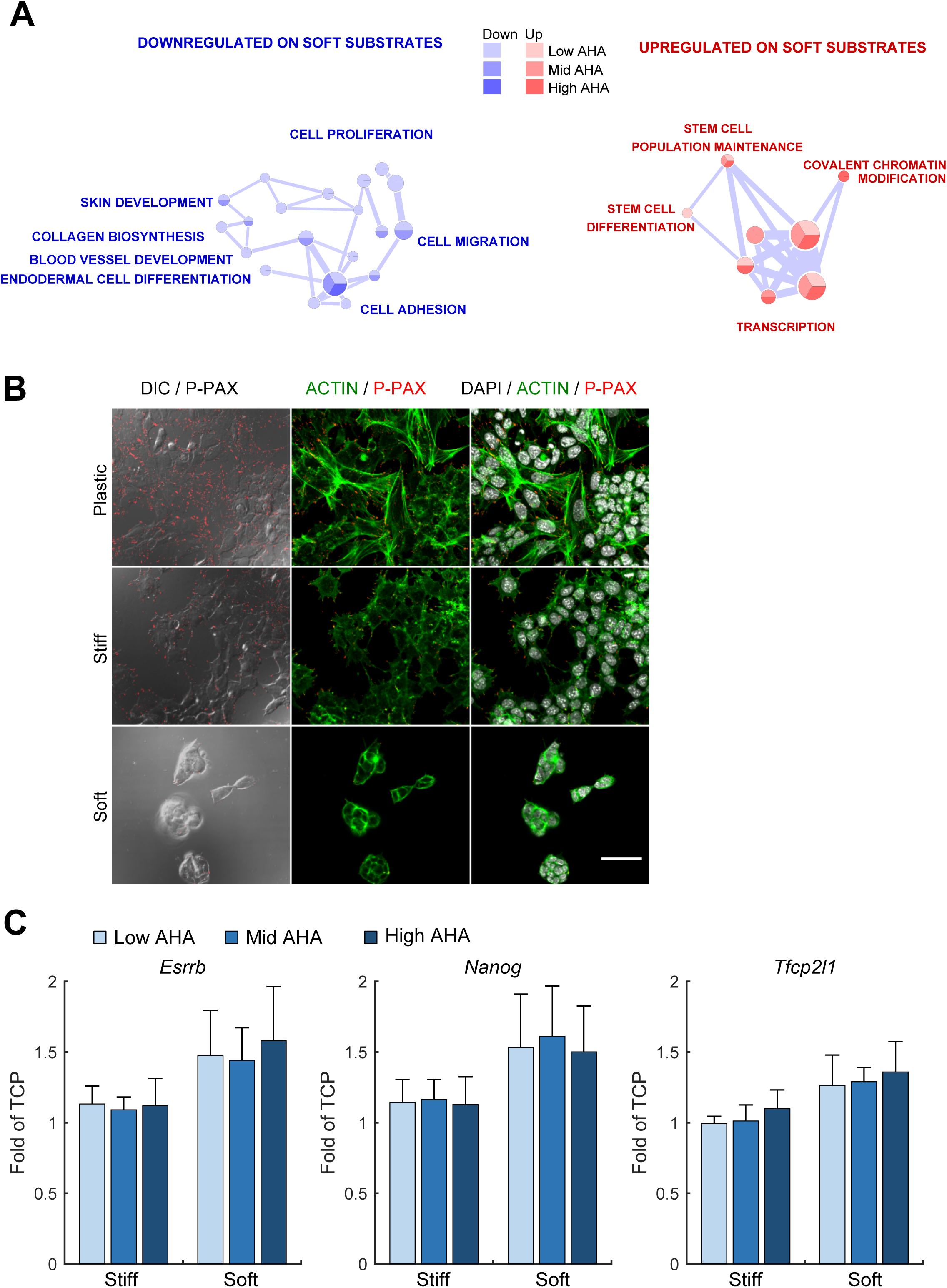
in relation to Figure 2. A – Network visualization of enriched biological processes due to substrate stiffness. In red are the processes enriched in upregulated genes, in blue the processes enriched in downregulated genes. Node size is proportional to the number of genes involved, and colour-charts inside each node indicate the adhesiveness for which the pathways were significantly enriched (padj < 0.1). Edge width indicates the similarity coefficient between nodes. B - Left panel: Overlay of phospho-paxillin (red) and DIC images. Central panel: Overlay of F-actin (Phalloidin – green) and phospho-paxillin (red). Right panel: Overlay of actin (green), phospho-paxillin (red) and DAPI (grey) for cells on tissue culture plastic (TCP), soft or stiff hydrogels. Focal adhesions marked by phospho-paxillin foci are only visible on TCP or stiff substrates. Scale bar: 50µm. C – Gene expression (mean +/− std) of naïve pluripotency genes *Esrrb, Nanog* and *Tfcpl21* for cells cultured for 24hr in S+L on fibronectin-coated TCP soft and stiff hydrogels and low AHA (light blue), mid AHA (cyan) and high AHA (dark blue) concentrations. Expressions were normalized to TCP (N = 3).

**Figure S3:**
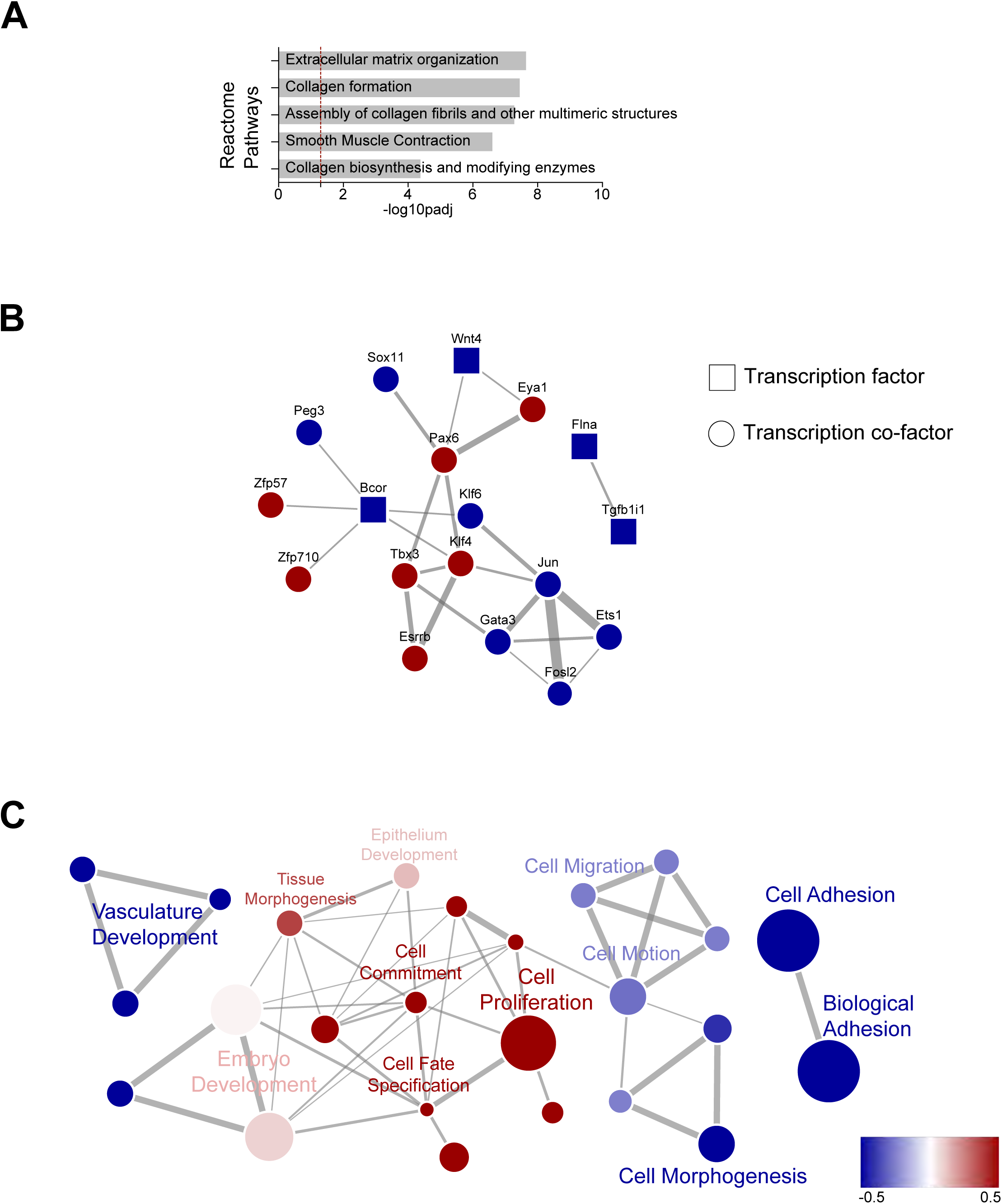
in relation to Figure 3, Gene Ontology analysis. A – Reactome pathways processes enriched for the bold genes in the centre of the Venn-diagram (n=219) of Figure 3A. B – Gene-Gene interaction network (https://string-db.org/) for the Transcription Factor (TF) and Transcription CoFactors (CoF) for the systematically modulated genes (n=219). Only the TF/CoF with at least 1 interaction are shown. Red / blue are for genes which are up -/ down-regulated on average. C – Top 25 enriched biological processes for genes up or downregulated in S-L conditions. The size is proportional to the number of genes that are enriched in the specific process. The colour is the z-score of the percentage of up/downregulated genes for each biological process.

**Figure S4:**
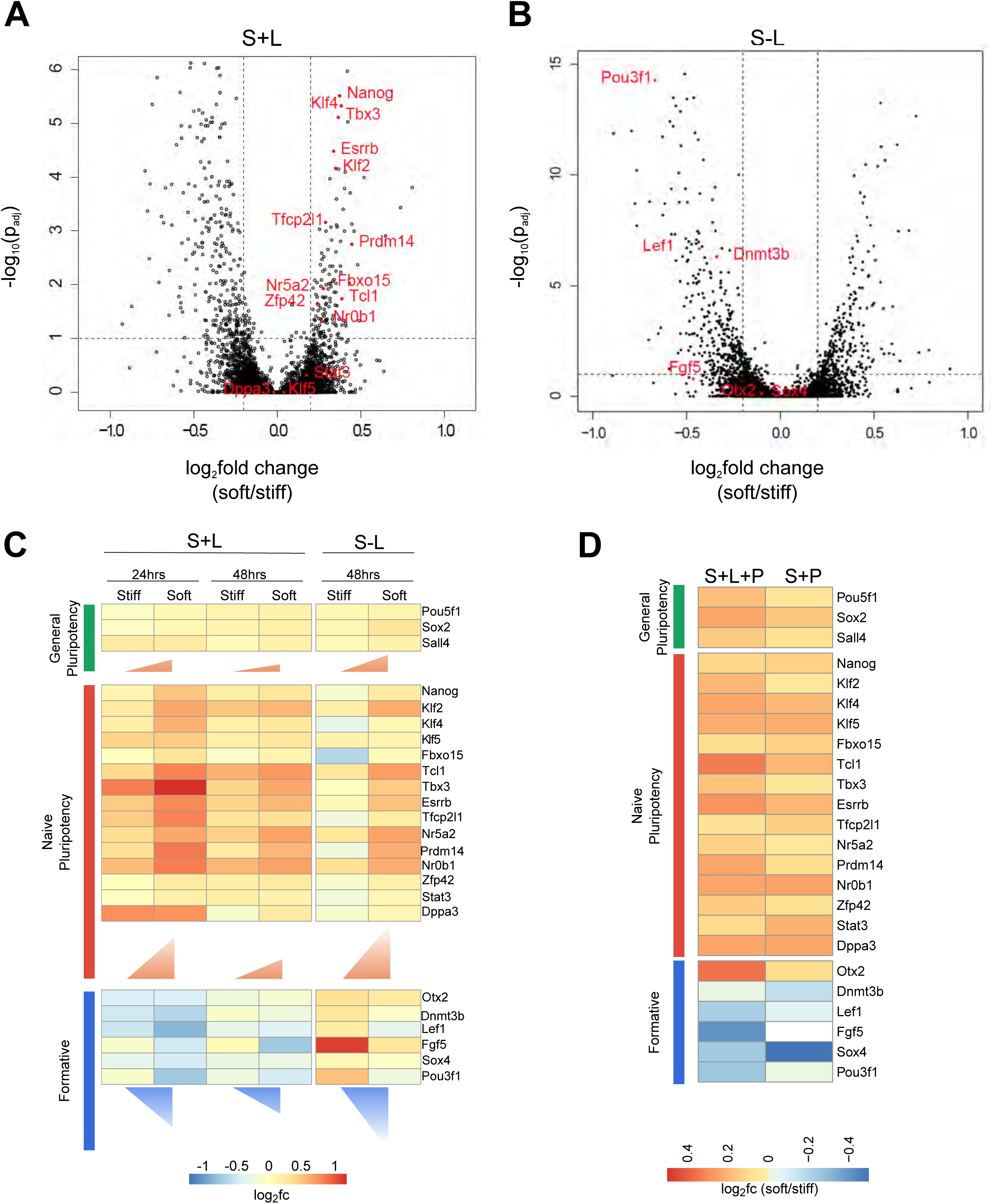
in relation to Figure 3C. A – Volcano plot showing significance (−log_10_(adjusted p-value)) versus fold change (log_2_(soft/stiff)) for the first condition shown in Figure 3C: S+L 24hrs. The dotted horizontal line shows the 10% false discovery rate level, any point above is significantly regulated. The vertical dotted lines indicate a log2 (soft/stiff) = +/− 0.2. The genes indicated in red are the naïve pluripotency markers shown in the heatmap of Figure 3C. B – Volcano plot showing significance (−log_10_(adjusted p-value)) versus fold change (log_2_(soft/stiff)) for the second condition shown in Figure 3C: S-L, 48hrs. The dotted horizontal line shows the 10% false discovery rate level, any point above is significantly regulated. The vertical dotted lines indicate a log2 (soft/stiff) = +/− 0.2. The genes indicated in red are the formative pluripotency markers shown in the heatmap of Figure 3C. C – Heatmap of log_2_fold change of expression for a selection of general pluripotency, naïve pluripotency and formative genes. All media conditions are compared to TCP control (S+L, 48hrs). The vertical side of each triangle is proportional to the average log_2_fc(soft/stiff) over all genes of a group for each condition D – Heatmap of log2(soft/stiff) for the same selection of genes as in Figure 3C and **S2D**. The conditions shown are serum+LIF+PD03 (“S+L+P”) and serum+PD03 (“S+P”).

**Figure S5:**
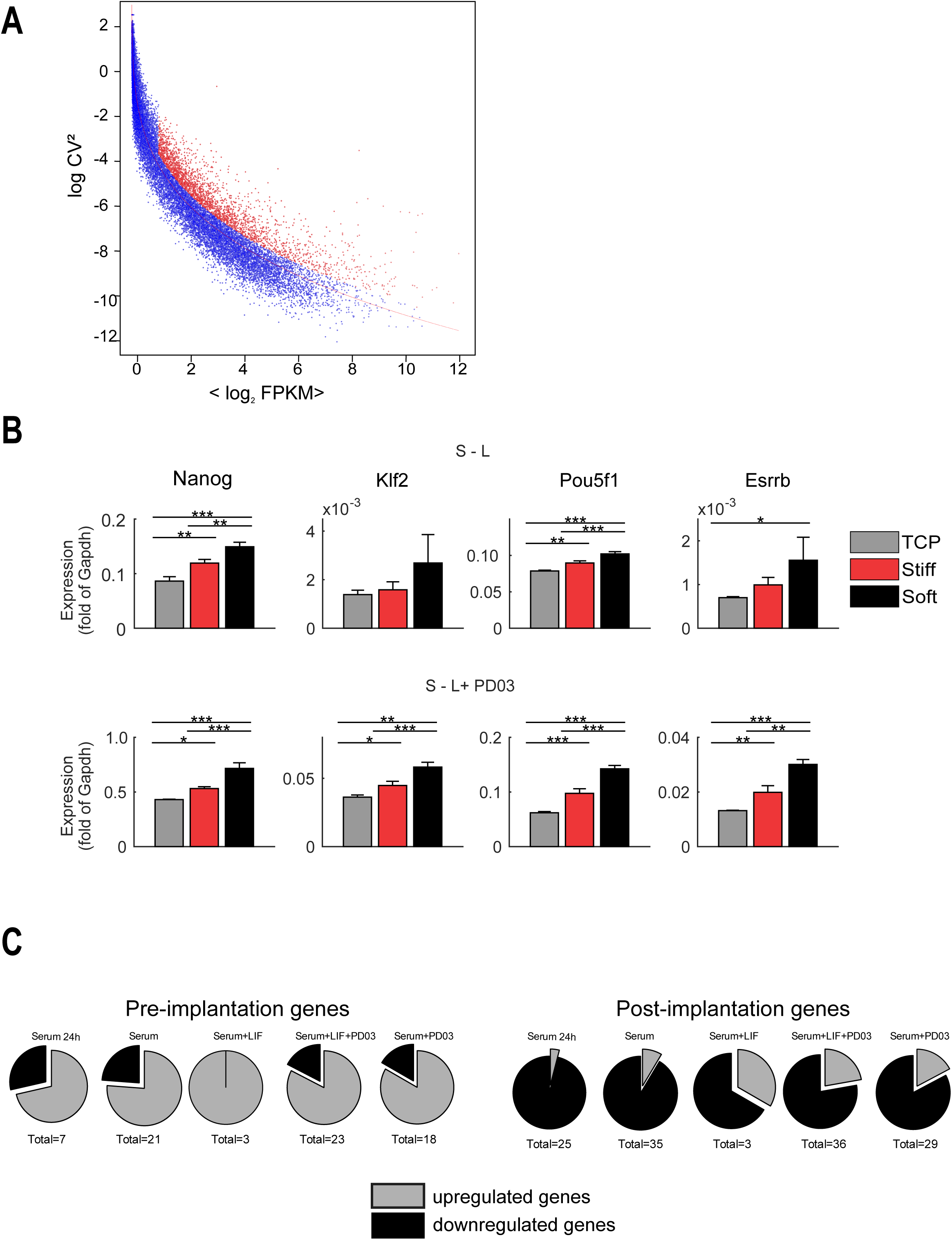
in relation to Figure 3D-E. A – Highly variable genes identified in S+L conditions (used for PCA plot in Figure 3D). The solid red line is a non-linear regression. Points in red are the genes that satisfy log_2_(FPKM) > 1 and with log(CV^2^) being 0.5 above the regression line (n=2579). B - Gene expression data for *Nanog, Klf2, Pou5f1 and Esrrb* for ES cells seeded on fibronectin-coated TCP (grey), stiff (red) and soft (black) hydrogels for 48 hrs. Cells were either initially cultured in S+L, then seeded at low density on the substrates in either S-L (serum without LIF), S-L+PD03. *Gapdh* was used as endogenous control (N=3). Error bars show standard deviation and *,**,*** indicate *p < 0.05, p < 0.01,* and *p < 0.001* respectively (ANOVA test). C - Proportion of significantly upregulated or downregulated genes on soft substrates compared to stiff substrates in each medium condition (in relation to Figure 3E).

**Figure S6:**
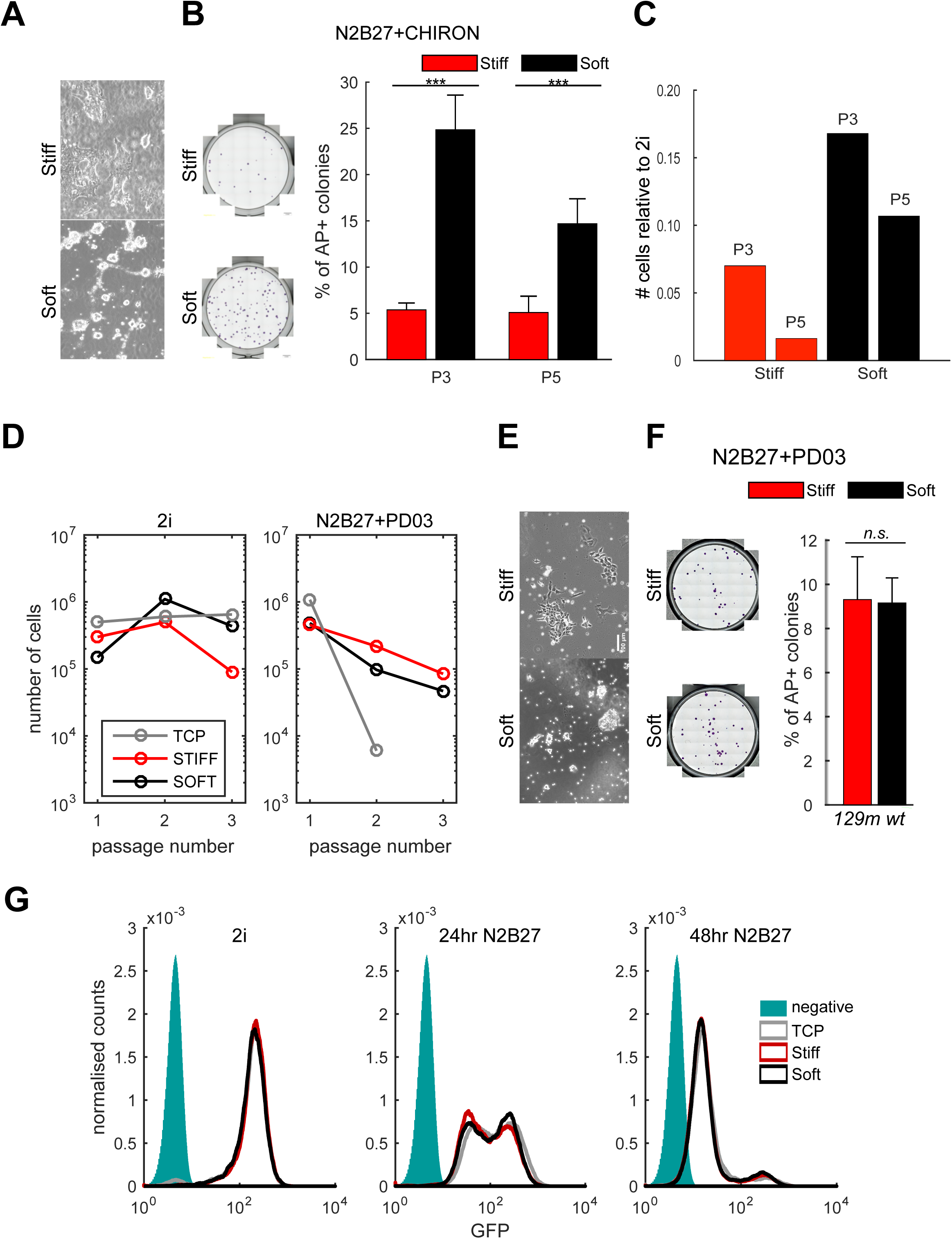
in relation to Figure 4. A – Brightfield images of cells on fibronectin-coated soft and stiff hydrogels after 3 passages in N2B27+CHIRON. B - Clonogenicity assay after cells were cultured in N2B27+CHIRON following protocol (ii) (see Figure 4A). (Left) Snapshots of staining for Alkaline Phosphatase (AP) after 3 passages. (Right) Quantification of the number of AP positive colonies as % of replated cells after 3 passages (P3) and 5 passages (P5) (N=8). C – Number of cells counted after 3 passages (P3) and 5 passages (P5) in N2B27+CHIRON on fibronectin-coated hydrogels. The cell numbers are given relative to the number of cells counted in the control conditions (2i). D – Number of cells counted at each passage in 2i, and N2B27+PD03, corresponding to **Figure S6F**. E – Brightfield images of cells on fibronectin-coated soft and stiff hydrogels after 3 passages in N2B27+PD03. F - Clonogenicity assay after cells were cultured in N2B27+PD03 following protocol (iii) (see Figure 4A). Cell line was from 129m background. (Left) Snapshots of staining for (AP) after 3 passages. (Right) Quantification of the number of AP positive colonies after 3 passages n % of replated cells (N=8). In all panels error bars show standard deviation and *,**,*** indicate *p < 0.05, p < 0.01,* and *p < 0.001* respectively (ANOVA test). G – Flow cytometry profiles of reporter line Rex1GFP::d2 cultured on TCP (grey), stiff (red) or soft (black) high AHA StemBond hydrogels for 0hr (left), 24hrs (centre) and 48hrs (right) in N2B27. The green filled histogram is a negative control. For each condition, histograms of two replicate samples were averaged and smoothed.

**Figure S7:**
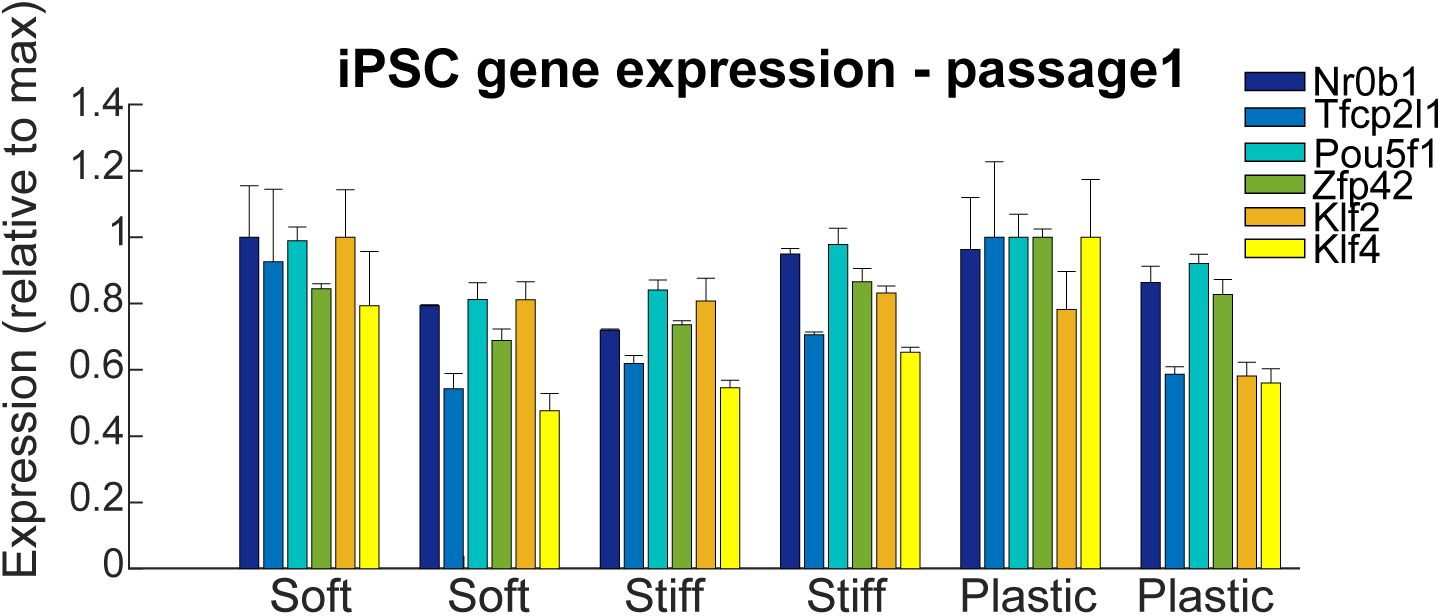
in relation to Figure 5. Expression (mean +/− std) of naïve pluripotency genes in iPSCs after 1 passage on TCP in 2i+LIF+bsd, following reprogramming on TCP, stiff or soft substrates. Expression for each replicate is presented relative to *Gapdh* then normalised to the highest value.

**Figure S8:**
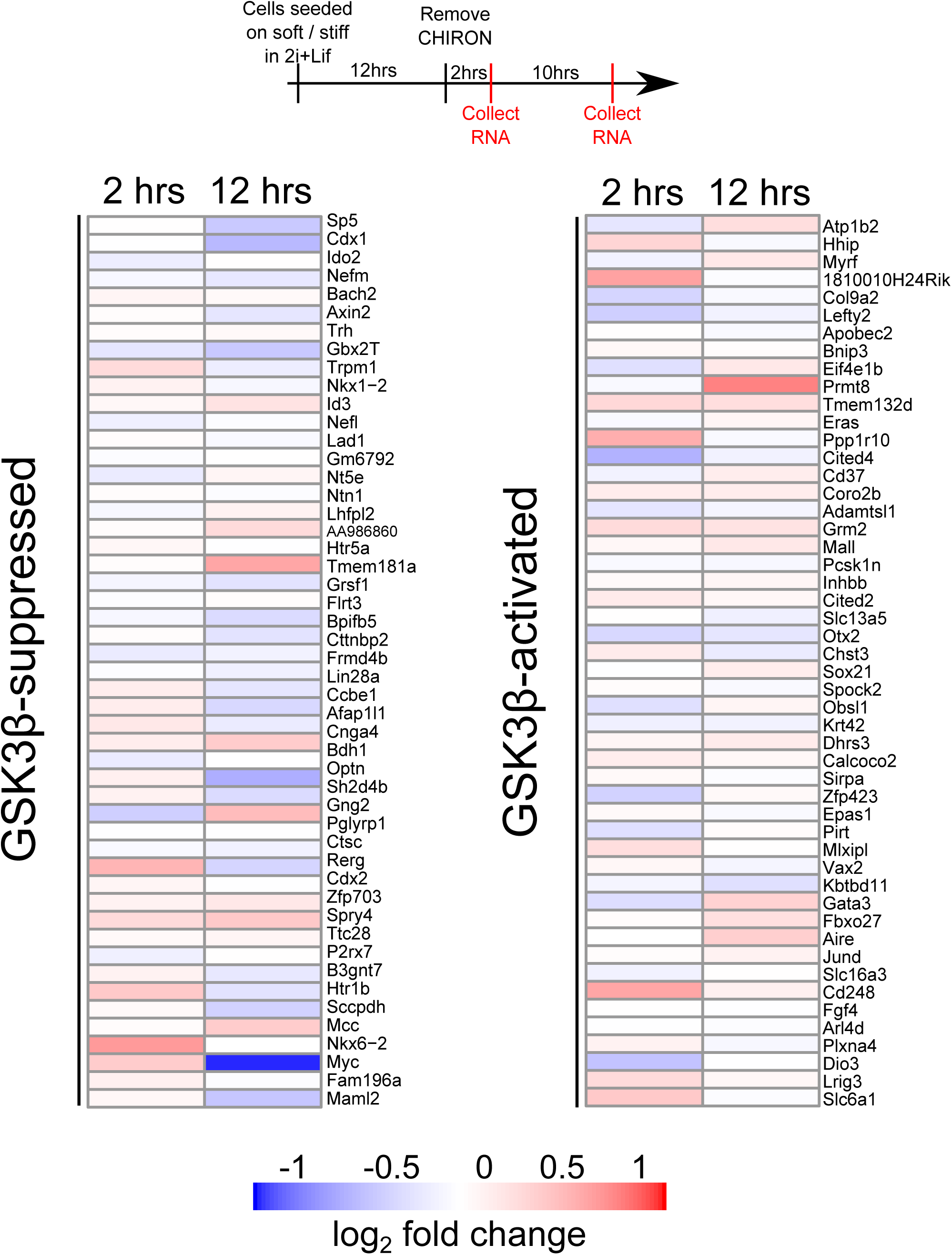
in relation to Figure 6, RNA sequencing after CHIRON removal. (Top) Design of experiments for RNA-sequencing indicating the timing of the different steps and CHIRON removal. Experiments performed in triplicate. (Bottom) Heatmaps of log_2_ fold change in expression between soft and stiff substrates after CHIRON removal. (Left) Top 50 activated genes and (right) Top 50 suppressed genes. The genes listed are those which are co-regulated after inhibitor removal on both soft and stiff and show the largest fold change on stiff substrates when compared to control in 2i+LIF.

